# Targeting glial PD-1/PD-L1 restores microglial homeostasis and attenuates neuronal hyperactivity in an Alzheimer’s disease model

**DOI:** 10.1101/2024.09.13.612389

**Authors:** Taeyoung Park, Leechung Chang, Seung Won Chung, Seowoo Lee, Sungjun Bae, Carlo Condello, Yong Ho Kim, Jaecheol Lee, Han-Joo Kim, Ho-Keun Kwon, Minah Suh

## Abstract

Alzheimer’s disease (AD) involves complex neuroimmune interactions. However, the role of immune checkpoint pathways in regulating the glial and neuronal functions of the AD brain remains unclear. This study aims to investigate how the brain-specific modulation of the PD-1/PD-L1 axis affects glial and neuronal function in an AD mouse model, which is characterized by the significant upregulation of microglial PD-1 and astrocytic PD-L1. A single intracortical administration of anti-PD-L1 antibody reshaped the glial microenvironment, which was accompanied by the restoration of impaired microglial process convergence. Astrocyte-specific *Pd-l1* knockdown using the pSico system validated the essential role of astrocytic PD-L1 in enhancing microglial responses. Notably, the blockade of the PD-1/PD-L1 signaling pathway increased the microglial P2RY12 expression, a key marker of microglial homeostasis, which likely contributed to the restoration of neuronal hyperactivity. Collectively, these findings confirmed the potential of targeting the astrocyte-microglia PD-1/PD-L1 axis to mitigate AD pathology.

**Teaser:** Modulating astrocytic PD-L1 restores microglial dynamics, enhances microglia-neuron crosstalk, and mitigates AD pathology.

## Introduction

AD is the leading cause of dementia and is characterized by the progressive accumulation of amyloid-beta (Aβ) plaques, neurofibrillary tangles, and chronic neuroinflammation, which results in memory and cognitive decline (*1*). Immunotherapy targeting the immune checkpoint pathway, particularly the programmed cell death protein 1/programmed death-ligand 1 (PD-1/PD-L1) axis, has emerged as a promising strategy for treating AD. Previous studies have suggested that the systemic administration of anti-PD-L1 antibody reduces Aβ burden and improves the cognitive outcomes in AD mouse models, primarily by activating peripheral immune responses and promoting the recruitment of monocyte-derived macrophages (MDMs) into the brain (*2, 3*). However, these studies have primarily focused on peripheral immune cells, leaving open questions about how the direct modulation of the PD-1/PD-L1 axis influences brain-resident cells, such as microglia and astrocytes, wherein increased PD-1/PD-L1 expression has been observed in both AD transgenic mice and human AD patients (*4*).

Unlike the systemic approaches, our study emphasizes the direct modulation of PD-1/PD-L1 signaling within the brain and its implications for glial and neuronal functions. Microglia, the brain’s intrinsic immune cells, are central to maintaining neuronal homeostasis; they monitor neuronal activities and modulate synaptic functions (*5–9*). In AD, microglia often exhibit a dysfunctional state characterized by impaired phagocytosis, reduced motility, and altered gene expression (*10–12*). Disruptions in these homeostatic functions may lead to an imbalance in excitatory and inhibitory signaling, which has been associated with neuronal hyperactivity, a hallmark of AD that contributes to network dysfunction and cognitive decline (*13–18*). While the relationship between neuronal hyperactivity in AD and microglial dysfunction remains complex, microglia are considered to play a key role in sensing and responding to neuronal activities through purinergic signaling, mediated by the P2RY12 receptor, subsequently regulating synaptic activity by suppressing glutamate release and receptor responsiveness (*6*). The expression of microglial P2RY12 is reduced in AD (*19–21*), suggesting its potential as a marker for disrupted microglial function in regulating hyperactive neurons. These findings suggest the need for strategies that not only enhance Aβ clearance but also restore microglial homeostatic functions, thereby facilitating microglia-neuron crosstalk in the AD brain.

Astrocytes also play a vital role in maintaining brain homeostasis by modulating neuronal activities, thereby regulating synaptic transmission and supporting microglial function (*22–27*). PD-L1 expressed on astrocytes has been implicated in neuroimmune interactions (*4, 28*), but its specific role in AD has not been fully investigated. Increased PD-1 and PD-L1 levels in microglia and astrocytes resemble their sustained expression in exhausted peripheral immune cells during chronic infections and tumors, wherein they help suppress excessive immune responses and prevent tissue damage (*29, 30*). In AD, such upregulation of PD-1/PD-L1 may similarly promote glial dysfunction by impairing immune surveillance and disrupting neuronal modulation. Consequently, targeting glial PD-1/PD-L1 signaling directly within the brain could provide a promising strategy to restore glial and neuronal function. Moreover, recent studies have suggested that modulating the brain intrinsic PD-1/PD-L1 pathway—for instance, by conditionally deleting PD-1 in hippocampal excitatory neurons in aged mouse models—improves synaptic transmission and cognitive function (*31*), emphasizing the significance of this pathway in regulating brain intrinsic interactions beyond peripheral immune regulation.

Although systemic PD-1/PD-L1 blockade has been shown to enhance peripheral immune responses in AD, its impact on brain-resident immune cells has not been determined (*2, 3*). Systemic anti-PD-L1 treatment promotes MDM infiltration into the brain via the choroid plexus, facilitating Aβ clearance (*3, 32*). However, the functional contribution of these infiltrating cells beyond phagocytosis remains unclear. The blood-brain barrier remains largely intact in several AD mouse models, restricting immune cell entry into the CNS infiltration (*33–35*), further emphasizing the need to understand the intrinsic roles of PD-1/PD-L1 signaling within the brain itself. While peripheral immune cells are increasingly recognized as key players in AD pathology interventions (*3, 36–40*), their interactions with brain-resident glia and neurons, as well as their influence on PD-1/PD-L1 signaling within the CNS, need to be fully investigated.

In this study, we investigated the effects of direct intracortical anti-PD-L1 administration on microglial function, neuronal activity, and Aβ pathology in a 5xFAD mouse model using real-time *in vivo* two-photon microscopy. Seven days after the intracortical injection, the anti-PD-L1 treatment effectively rescued microglial process motility and restored neuronal baseline calcium activity, while significantly reducing the Aβ plaques and increasing plaque-associated microglia. Furthermore, astrocyte-specific *Pd-l1* knockdown showed effects similar to those resulting from the inhibition of the glial PD-1/PD-L1 pathway. In general, our findings suggest that brain intrinsic glial PD-1/PD-L1 modulation reduces Aβ pathology along with restoring functional interactions among microglia, astrocytes, and neurons. This work is consistent with previous studies that focused on peripheral immune mechanisms and determined the brain-specific mechanisms underlying immune checkpoint modulation. By clarifying the mechanistic basis of glial-neuron interactions in AD, our findings pave the way for targeted approaches aimed at restoring brain homeostasis and mitigating neurodegeneration.

## Results

### Intracortical injection of the anti-PD-L1 antibody increases microglial process convergence in 5xFAD mice

To investigate the expression of PD-1 and PD-L1 in AD pathology, their levels in the cortex and subiculum of 5xFAD and age-matched wild-type (WT) mice were analyzed. The microglia in 5xFAD mice revealed a significantly increased PD-1 expression, whereas astrocytes exhibited increased PD-L1 expression with an age-dependent rise correlating with disease progression (fig. S1). The neurons had lower PD-1 and PD-L1 levels than the glial cells (Fig. S1, E, and H), and the colocalization analysis confirmed that microglial PD-1 and astrocytic PD-L1 expression aligned with the markers of AD pathology.

**Figure 1.**
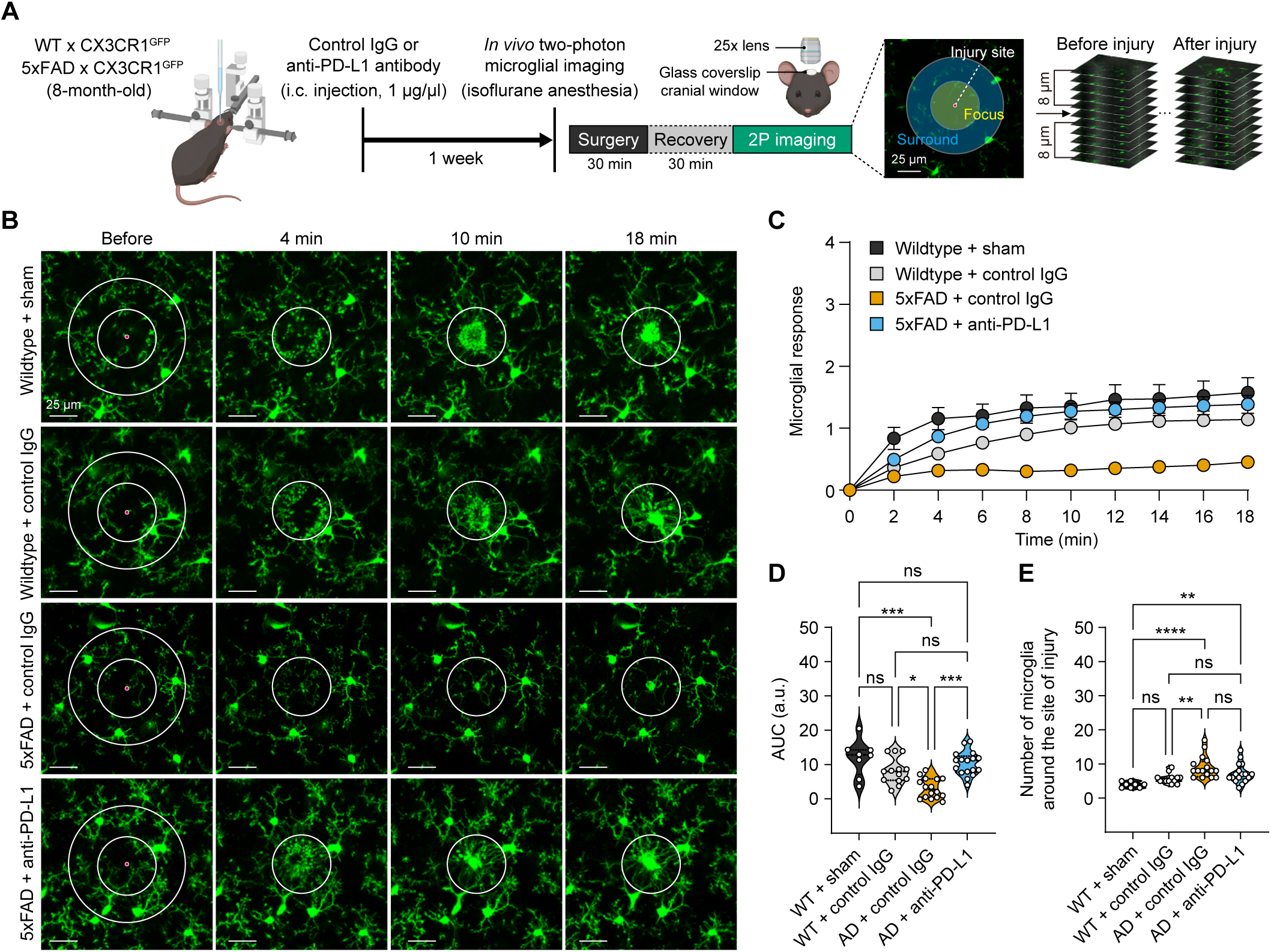
Blockade of PD-1/PD-L1 signaling in the brain increases microglial process convergence in response to local brain injury in 5xFAD mice. (**A**) The experimental scheme of *in vivo* two-photon (2P) microglial time-lapse imaging with focal laser injury seven days after a single intracortical injection of antibody in WT/CX-3CR1^GFP^ or 5xFAD/CX3CR1^GFP^ mice by crossing 5xFAD with CX3CR1^GFP/GFP^ mice. (**B**) Representative time-lapse imaging of laser injury-targeted microglial process conver-gence. (**C**) Time course of microglial process kinetics toward a laser-induced lesion (WT + sham; *n* = 8, WT + control IgG; *n* = 13, 5xFAD + control IgG; *n* = 16, 5xFAD + anti-PD-L1; *n* = 17). Microglial responses = If(t)-If(0)/Is(t). If indicates the intensity of the focus, whereas Is indicates the intensity of the surrounding area (mean ± s.e.m.). (**D**) The area under the curve (AUC) of the accumulated graphs in (C) (Kruskal-Wallis test with Dunn’s post-hoc comparisons test; WT + sham vs. WT + control IgG; *p* = 0.8985, WT + sham vs. AD + control IgG; \*\*\**p* = 0.0005, WT + sham vs. AD + anti-PD-L1; *p* = 1.000, WT + control IgG vs. AD + control IgG; \**p* = 0.0288, WT + control IgG vs. AD + anti-PD-L1; *p* = 1.000, AD + control IgG vs. AD + anti-PD-L1; \*\*\**p* = 0.0001, ns, not significant). (**E**) The number of microglia around the site of focal injury within 100 μm (Kruskal-Wallis test with Dunn’s post-hoc comparisons test; WT + sham vs. WT + control IgG; *p* = 0.2956, WT + sham vs. AD + control IgG; \*\*\*\**p* < 0.0001, WT + sham vs. AD + anti-PD-L1; \*\**p* = 0.0039, WT + control IgG vs. AD + control IgG; \*\**p* = 0.0096, WT + control IgG vs. AD + anti-PD-L1; *p* = 0.7014, AD + control IgG vs. AD + anti-PD-L1; *p* = 0.5063, ns, not significant).

We also focused on the specific effects of blocking the PD-1/PD-L1 pathway on microglial homeostatic function, utilizing real-time *in vivo* two-photon imaging. To achieve this, we administered an anti-PD-L1 antibody directly into the cortical region of 8-month-old 5xFAD mice (Fig. 1A). To enable the visualization of microglia in the AD brain, fluorescently labeled microglia were generated by crossbreeding 5xFAD mice with CX3CR1-green fluorescent protein (GFP) mice. Following a focal laser brain tissue injury to induce ATP release, we observed a microglial process convergence to the injury site in the cortex. The 5xFAD mice treated with control IgG revealed a significant reduction in microglial process convergence compared with WT mice, but this impairment was significantly rescued by intracortical anti-PD-L1 treatment (Fig. 1, B to D). Because microglial behavior can be influenced by cell density near the injury site, we confirmed that the microglial numbers in the imaging area did not differ between the control IgG- and anti-PD-L1-treated 5xFAD groups (Fig. 1E). These findings emphasize that the homeostatic function of plaque-associated microglia is specifically restored through intracortical anti-PD-L1 injection, suggesting the importance of localized treatment strategies for effective intervention.

To determine whether the systemic administration of anti-PD-L1 could produce similar effects, we performed intraperitoneal (i.p.) injections of the antibody in 5xFAD mice, delivering a single dose (0.5 mg per mouse) (*3*). Unlike the intracortical approach, the i.p. administration of anti-PD-L1 failed to restore microglial process convergence, suggesting the necessity of direct brain-targeted delivery for the effective modulation of microglial function (Fig. S2).

### Astrocyte-specific *Pd-l1* knockdown increases the microglial process convergence in 5xFAD mice

To determine the cell-specific role of PD-L1 (encoded by the *Cd274* gene) in restoring microglial homeostatic activity, the pSico (the plasmid for stable RNAi, conditional) system (*41*) was used to selectively knock down *Pd-l1* in astrocytes or neurons. Astrocytic *Pd-l1* knockdown was achieved using pSico-*Cd274* shRNA with GFAP-Cre, whereas neuronal *Pd-l1* knockdown was performed using pSico-*Cd274* shRNA with CaMK2a-Cre (Fig. 2, A and B). The knockdown of *Pd-l1* in astrocytes significantly restored microglial process convergence in 5xFAD mice, closely mirroring the effects of intracortical anti-PD-L1 antibody injection (Fig. 2, C, E, and F). In contrast, the neuronal *Pd-l1* knockdown failed to rescue the impaired microglial process convergence in 5xFAD mice (Fig. 2, D, G, and H). These findings suggest the essential role of astrocytic PD-L1 in regulating the microglial process dynamics and that the restorative effects of PD-L1 blockade are predominantly mediated through astrocyte-microglia crosstalk.

**Figure 2.**
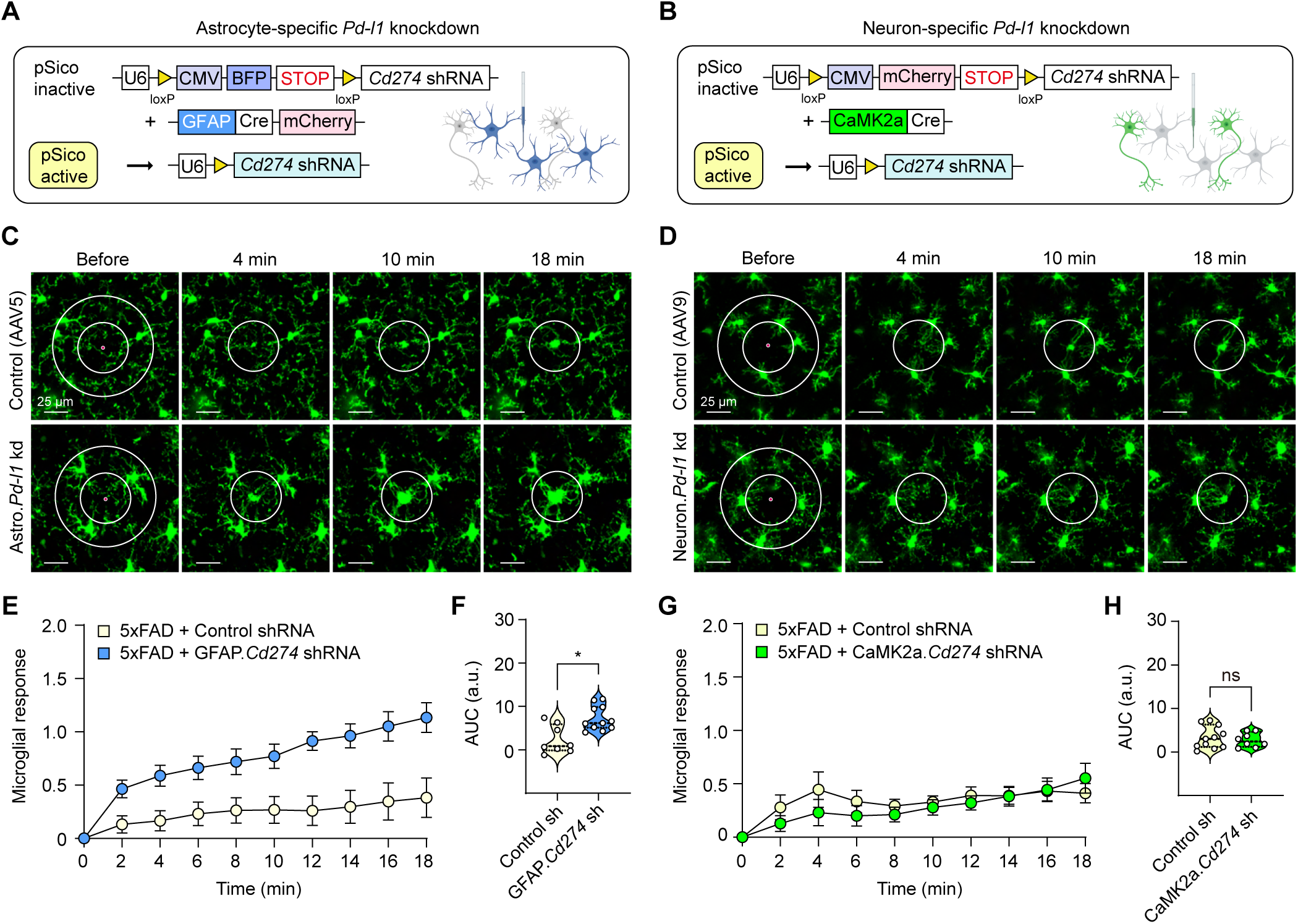
Astrocyte-specific Pd-l1 knockdown increases microglial process convergence in response to local brain injury in 5xFAD mice. (**A**) The experimental scheme of *in vivo* microglial time-lapse imaging with focal laser injury two weeks after the intracortical injection of the control virus (a mixture of AAV5-GFAP-Cre-mCherry and AAV5-pSico-scrambled shRNA-BFP) or astrocyte-specific *Pd-l1* knockdown virus (a mixture of AAV5-GFAP-Cre-mCherry and AAV5-pSico-Cd274 shRNA-BFP) in 5xFAD/CX-3CR1^GFP^ mice. (**B**) The experimental scheme of *in vivo* microglial time-lapse imaging with focal laser injury two weeks after the intracortical injection of the control virus (a mixture of AAV9-CaMK2a-Cre and AAV9-pSico-scrambled shRNA-mCherry) or excitatory neuron-specific *Pd-l1* knockdown virus (a mixture of AAV9-CaMK2a-Cre and AAV9-pSico-Cd274 shRNA-mCherry) in 5xFAD/CX3CR1^GFP^ mice. (**C** and **D**) Representative time-lapse imaging of laser injury-targeted microglial process convergence. (**E**) Time course of microglial process kinetics toward a laser-induced lesion (5xFAD + control shRNA; *n* = 8, 5xFAD + GFAP.Cd274 shRNA; *n* = 11, mean ± s.e.m.). (**F**) The AUC of the accumulated graphs in (E) (Mann–Whitney U test, \**p* = 0.017). (**G**) Time course of microglial process kinetics toward a laser-induced lesion (5xFAD + control shRNA; *n* = 11, 5xFAD +CaMK2a.Cd274 shRNA; *n* = 8, mean ± s.e.m.). (**H**) The AUC of the accumulated graphs in (G) (Mann–Whitney U test; *p* = 0.869, ns, not significant).

### Intracortical injection of anti-PD-L1 antibody increased microglial P2RY12 expression and restored abnormal spontaneous neuronal hyperexcitability in 5xFAD mice

To assess the effects of anti-PD-L1 treatment on microglial gene expression, P2RY12, a critical marker of microglial homeostasis and a regulator of neuronal activity, was evaluated. The P2RY12 levels were significantly reduced in 5xFAD mice, particularly in plaque-associated microglia, but were restored around plaques following anti-PD-L1 treatment (Fig. 3, A to C). Given that P2RY12 enables microglia to sense extracellular ATP/ADP and regulate neuronal activity, its restoration is essential for mitigating neuronal hyperactivity.

**Figure 3.**
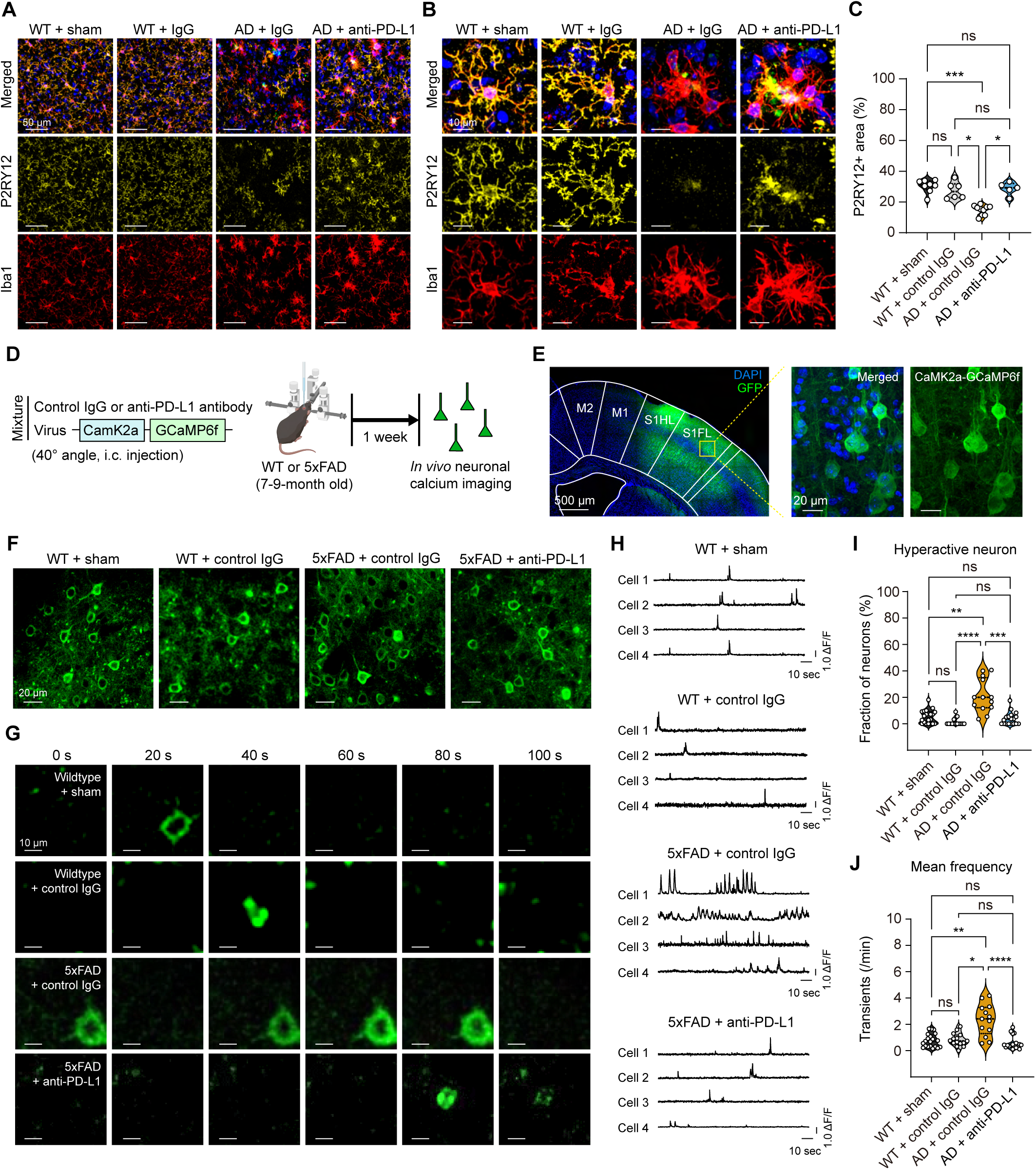
Blockade of PD-1/PD-L1 signaling in the brain rescues P2RY12 expression and neuronal activity in 5xFAD mice. (**A**) Representative images of P2RY12^+^ (yellow) and Iba1^+^ (red) microglia seven days after a single intracortical injection. (**B**) Magnification images in (A). (**C**) The quantitation of the P2RY12-positive area (WT + sham; *n* = 8 mice, WT + control IgG; *n* = 6 mice, 5xFAD + control IgG; *n* = 10 mice, 5xFAD + anti-PD-L1; *n* = 5 mice, Kruskal-Wallis test with Dunn’s post-hoc comparisons test; WT + sham vs. WT + control IgG; *p* = 1.000, WT + sham vs. AD + control IgG; \*\*\**p* = 0.0003, WT + sham vs. AD + anti-PD-L1; *p* = 1.000, WT + control IgG vs. AD + control IgG; \**p* = 0.0211, WT + control IgG vs. AD + anti-PD-L1; *p* = 1.000, AD + control IgG vs. AD + anti-PD-L1; \**p* = 0.0198, ns, not significant). (**D**) The experimental scheme of *in vivo* neuronal calcium imaging seven days after the single intracortical injection of the mixture of antibody and CaMK2a-GCaMP6f virus in 5xFAD mice. (**E**) The confocal images showing the expression of the injected virus (green, CaMK2a-GCaMP6f; blue, DAPI) in the S1FL. (**F**) Representative *in vivo* two-photon fluorescence images of GCaMP6f-ex-pressing cortical neurons from layer II/III of the control IgG- or anti-PD-L1 antibody-injected 7- to 9-month-old 5xFAD mice and age-matched WT mice. (**G**) The fluorescence images of the spontaneous calcium transients of a single excitatory neuron in each group. (**H**) The representative calcium fluorescent traces of neurons. (**I**) The fractions of the hyperactive neurons in the sham or control IgG-injected WT mice and control IgG- or anti-PD-L1-injected AD mice (WT + sham; *n* = 20, WT + control IgG; *n* = 15, 5xFAD + control IgG; *n* = 13, 5xFAD + anti-PD-L1; *n* = 18, Kruskal-Wallis test with Dunn’s post-hoc comparisons test; WT + sham vs. WT + control IgG; *p* = 0.1790, WT + sham vs. AD + control IgG; \*\**p* = 0.0039, WT + sham vs. AD + anti-PD-L1; *p* = 1.000, WT + control IgG vs. AD + control IgG; \*\*\*\**p* < 0.0001, WT + control IgG vs. AD + anti-PD-L1; *p* = 1.000, AD + control IgG vs. AD + anti-PD-L1; \*\*\**p* = 0.0002, ns, not significant). (**J**) The mean frequency of neuronal calcium transients (Kruskal-Wallis test with Dunn’s post-hoc comparisons test; WT + sham vs. WT + control IgG; *p* = 1.0000, WT + sham vs. AD + control IgG; \*\**p* = 0.0015, WT + sham vs. AD + anti-PD-L1; *p* = 1.000, WT + control IgG vs. AD + control IgG; \**p* = 0.0200, WT + control IgG vs. AD + anti-PD-L1; *p* = 0.9558, AD + control IgG vs. AD + anti-PD-L1; \*\*\*\**p* < 0.0001, ns, not significant).

Subsequently, to directly investigate the impact of PD-1/PD-L1 blockade on microglia-neuron crosstalk, *in vivo* two-photon calcium imaging was used to monitor the excitatory neuronal activity. By co-delivering AAV9-CaMK2a-GCaMP6f with the anti-PD-L1 antibody into the cortex of 8-month-old 5xFAD mice, the neuronal calcium transients were visualized at the single-cell level (Fig. 3 D to F). Anti-PD-L1 treatment significantly reduced the frequency and fraction of hyperactive neurons in 5xFAD mice, suggesting a recovery of neuronal activity toward normal levels (Fig. 3, G to J). These results emphasize the role of PD-L1 blockade in restoring microglial homeostasis and reshaping microglia-neuron interactions, thereby reducing a hallmark of AD pathology.

### Intracortical injection of anti-PD-L1 antibody restores sensory-evoked neurovascular responses and neural activity in 5xFAD mice

To assess the effects of anti-PD-L1 administration on sensory-evoked neurovascular responses and neural activity in the primary somatosensory forelimb cortex (S1FL), we performed *in vivo* intrinsic optical signal (IOS) imaging to assess the cerebral blood volume (CBV) changes and local field potential (LFP) recordings to monitor neuronal ensemble activity during 20 s of electrical forepaw stimulation (Fig. 4, A and B). 5xFAD mice treated with control IgG exhibited significant reduction of evoked CBV changes, including diminished peak amplitude and AUC, compared with the WT controls (Fig. 4, C, and D). However, intracortical anti-PD-L1 administration rescued this impairment, restoring the CBV changes to levels comparable to those in WT mice. The onset time of CBV changes showed no differences between the groups. Additionally, evoked neural responses, measured by relative increases in LFP power during stimulation, were significantly reduced in 5xFAD mice but significantly improved following anti-PD-L1 administration (Fig. 4, E to G). These results revealed that anti-PD-L1 treatment restores neurovascular coupling and emphasizes the significance of glial PD-1/PD-L1 signaling in maintaining proper neurovascular and neural function in the brain.

**Figure 4.**
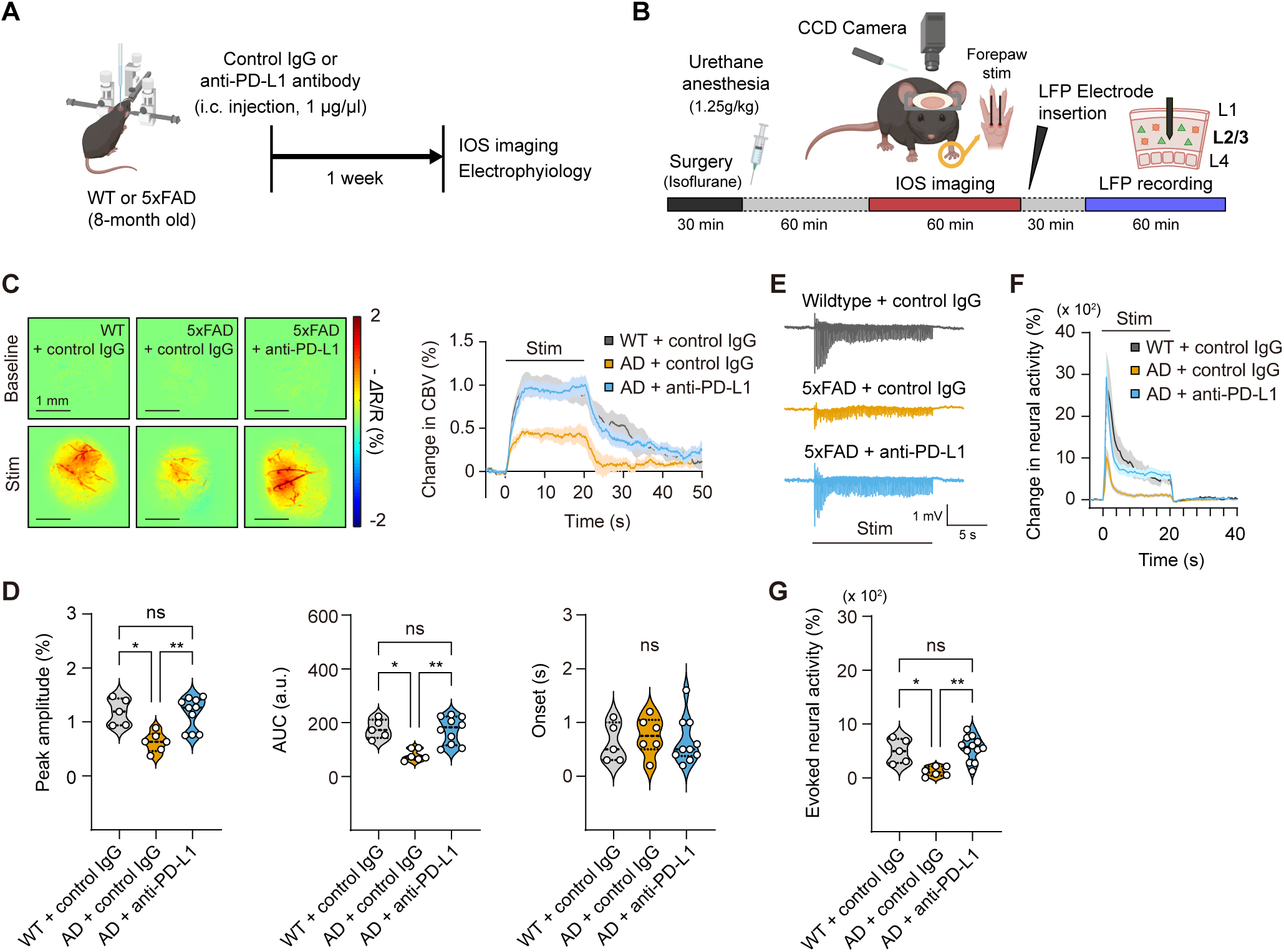
Blockade of PD-1/PD-L1 signaling in the brain enhances neurovascular function in 5xFAD mice. (**A**) The experimental scheme of *in vivo* neurovascular imaging and electrophysiological recording seven days after a single intracortical injection of antibody in the WT or 5xFAD mice. (**B**) The experimental procedures for stereotaxic surgery, IOS imaging, and LFP recording to assess the changes in the CBV and neural activity in the primary somatosensory cortex of the forelimb region (S1FL) during electrical forepaw stimulation. (**C**) Representative spatiotemporal IOS images and time course traces of the relative changes in the evoked CBV signal following forepaw stimulation (20 seconds), relative to the pre-stimulus baseline levels (WT + control IgG; *n* = 5, 5xFAD + control IgG; *n* = 6, 5xFAD + anti-PD-L1; *n* = 10). (**D**) The properties of the evoked CBV signal including the maximum evoked change, AUC, and time of onset (Kruskal-Wallis test with Dunn’s post-hoc comparisons test; Peak amplitude; WT + control IgG vs. AD + control IgG; \**p* = 0.0169, WT + control IgG vs. AD + anti-PD-L1; *p* = 0.9000, AD + control IgG vs. AD + anti-PD-L1; \*\**p* = 0.0091, AUC; WT + control IgG vs. AD + control IgG; \*\**p* = 0.0266, WT + control IgG vs. AD + anti-PD-L1; *p* = 1.000, AD + control IgG vs. AD + anti-PD-L1; \*\**p* = 0.0097, Onset; no significant difference was detected among the groups by the Kruskal-Wallis test, ns, not significant). (**E**) Representative raw LFP traces. (**F**) Time course traces of the relative changes in the evoked neural activity following forepaw stimulation (20 seconds), relative to the pre-stimulus baseline levels (WT + control IgG; *n* = 5, 5xFAD + control IgG; *n* = 6, 5xFAD + anti-PD-L1; n = 12). (**G**) Evoked LFP responses within the frequency range of 2–100 Hz (Kruskal-Wallis test with Dunn’s post-hoc comparisons test; WT + control IgG vs. AD + control IgG; \**p* = 0.0390, WT + control IgG vs. AD + anti-PD-L1; *p* = 1.000, AD + control IgG vs. AD + anti-PD-L1; \*\**p* = 0.0032, ns, not significant).

### Intracortical injection of anti-PD-L1 antibody promotes the microglial phagocytosis of Aβ in 5xFAD mice

The effects of anti-PD-L1 intracortical injection on the clearance of Aβ plaques in 5xFAD mice were investigated. Isotype IgG was injected into the left hemisphere of the cortex as a control, while the right hemisphere of the cortex received anti-PD-L1 administration in 5xFAD mice. Seven days after anti-PD-L1 injection, the size of the Aβ plaques in the injection areas was evaluated (Fig. 5A). A shift toward smaller Aβ plaques in the anti-PD-L1-injected area compared with the control IgG-injected area was observed (Fig. 5B). Of note, our findings revealed a significant decrease in the size of Aβ plaques compared with the control IgG, suggesting an augmented clearance of Aβ plaques facilitated by anti-PD-L1 intracortical injection (Fig. 5C).

**Figure 5.**
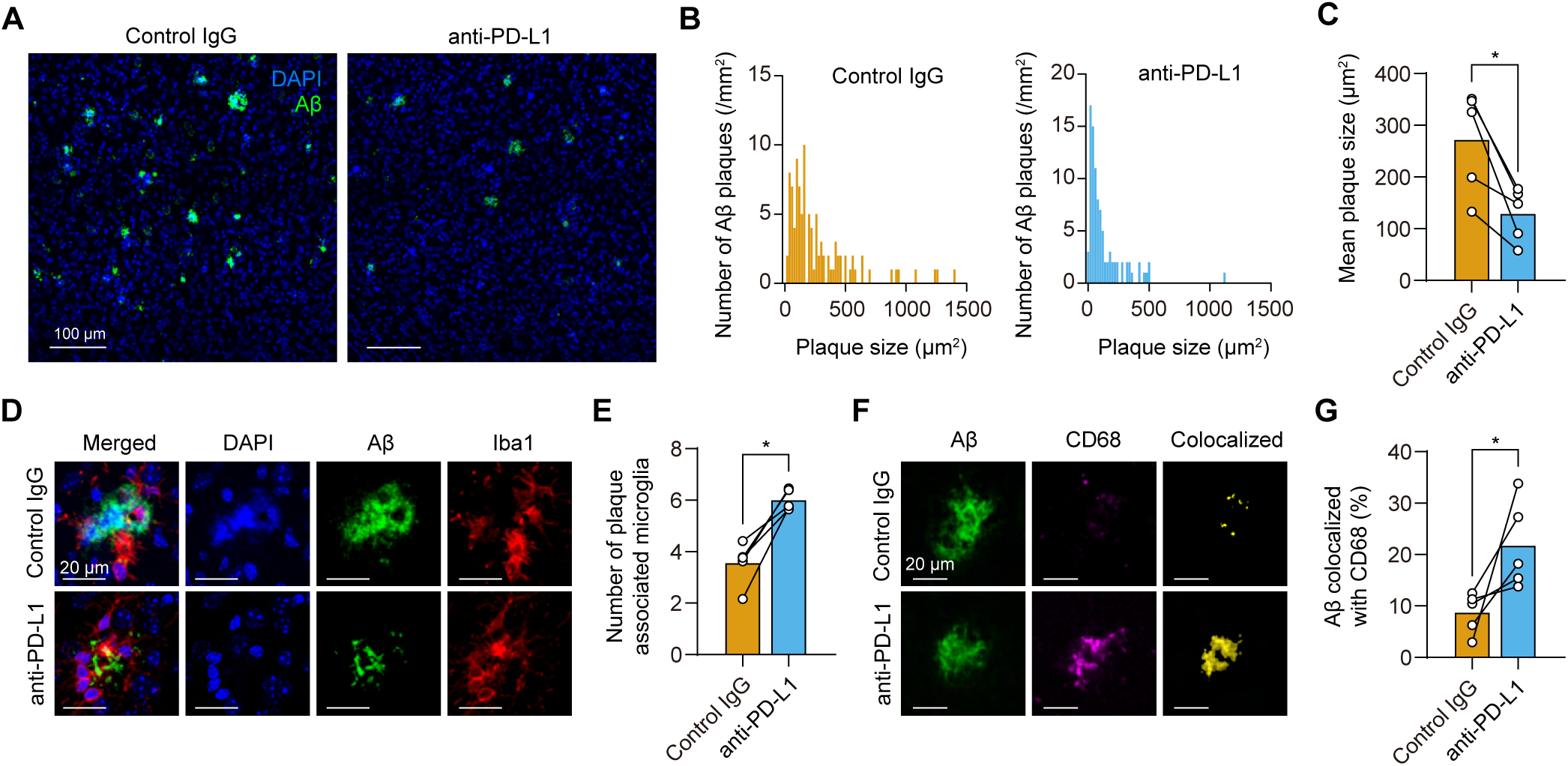
Blockade of PD-1/PD-L1 signaling in the brain improves Aβ pathology and promotes microglial phagocytosis in 5xFAD mice. (**A**) Representative images of the control IgG- or anti-PD-L1-injected cortex of 8-month-old 5xFAD mice stained with DAPI (blue) for nuclei and 6E10 (green) for Aβ, which were assessed seven days after a single intracortical injection. (**B**) The quantitation of the number of Aβ plaques with different sizes (control IgG group, 101 plaques/5 mice; anti-PD-L1 group, 93 plaques/5 mice). (**C**) The mean Aβ plaque size in (B) (*n* = 5 mice, Wilcoxon signed-rank test; \**p* = 0.043). (**D**) Representative images of antibody-injected 5xFAD mice stained with DAPI (blue) for nuclei, 6E10 (green) for Aβ, and Iba1 (red) for microglia. (**E**) The quantitation of the number of plaque-associated microglia (*n* = 5 mice, Wilcoxon signed-rank test; \**p* = 0.043). (**F**) Representative images of the antibody-injected cortex of 5xFAD mice stained with 6E10 (green) for Aβ and CD68 (magenta) for the phagocytic phenotype of microglia. Aβ colocalized with CD68 was shown. To assess the colocalization of an Aβ plaque and CD68, z-stacks were acquired and 3D colocalization analysis was conducted using the IMARIS software (Bitplane). (**G**) The quantitative assessment of the percentage area of the colocalization of Aβ with CD68. Plaques above 300 μm^2^ in size were selected for analysis (*n* = 5 mice, Wilcoxon signed-rank test; \**p* = 0.043).

To confirm that the reduction in Aβ plaque size occurred through microglia-mediated mechanisms, the number of plaque-associated microglia, a subtype of microglial cells that are found near the Aβ plaques in brain tissue, was evaluated (*42, 43*). An increase in the amount of plaque-associated microglia was observed in the cortex after anti-PD-L1 injection (Fig. 5, D, and E). The quantitative analysis of the colocalization area between CD68 and Aβ, indicating active microglial engulfment, phagocytosis, and the clearance of Aβ, was performed in the regions injected with control IgG or anti-PD-L1. Our investigations revealed a significant increase in the colocalization area of CD68 with the Aβ plaques following anti-PD-L1 injection (Fig. 5, F and G), which suggests an enhancement in the microglial lysosomal activity. Thus, these results revealed that anti-PD-L1 administration facilitates the efficient microglial clearance of the Aβ plaques in the brain, thereby contributing to the therapeutic efficacy in AD.

### Peripheral immune cell infiltration is induced by anti-PD-L1 antibody administration into the brain

To specifically investigate the impact of anti-PD-L1 on the brain, the antibody was injected into the primary somatosensory cortex of the forelimb region (S1FL) in 5xFAD mice, assessing both the antibody distribution and target cell identification (Fig. 6A). To rule out the potential contralateral effects, we confirmed that antibody diffusion remained localized to the injection site, as demonstrated by immunostaining, with no observed cross-hemispheric spread, indicating a highly confined impact. One day after the injection, immunostaining targeting the rat IgG revealed that the injected anti-PD-L1 was well distributed over the cortical region (approximately 1–1.5 mm³ in volume), with a particularly strong binding to the astrocytes expressing GFAP compared with the control IgG-injected area (Fig. 6B, fig. S3).

**Figure 6.**
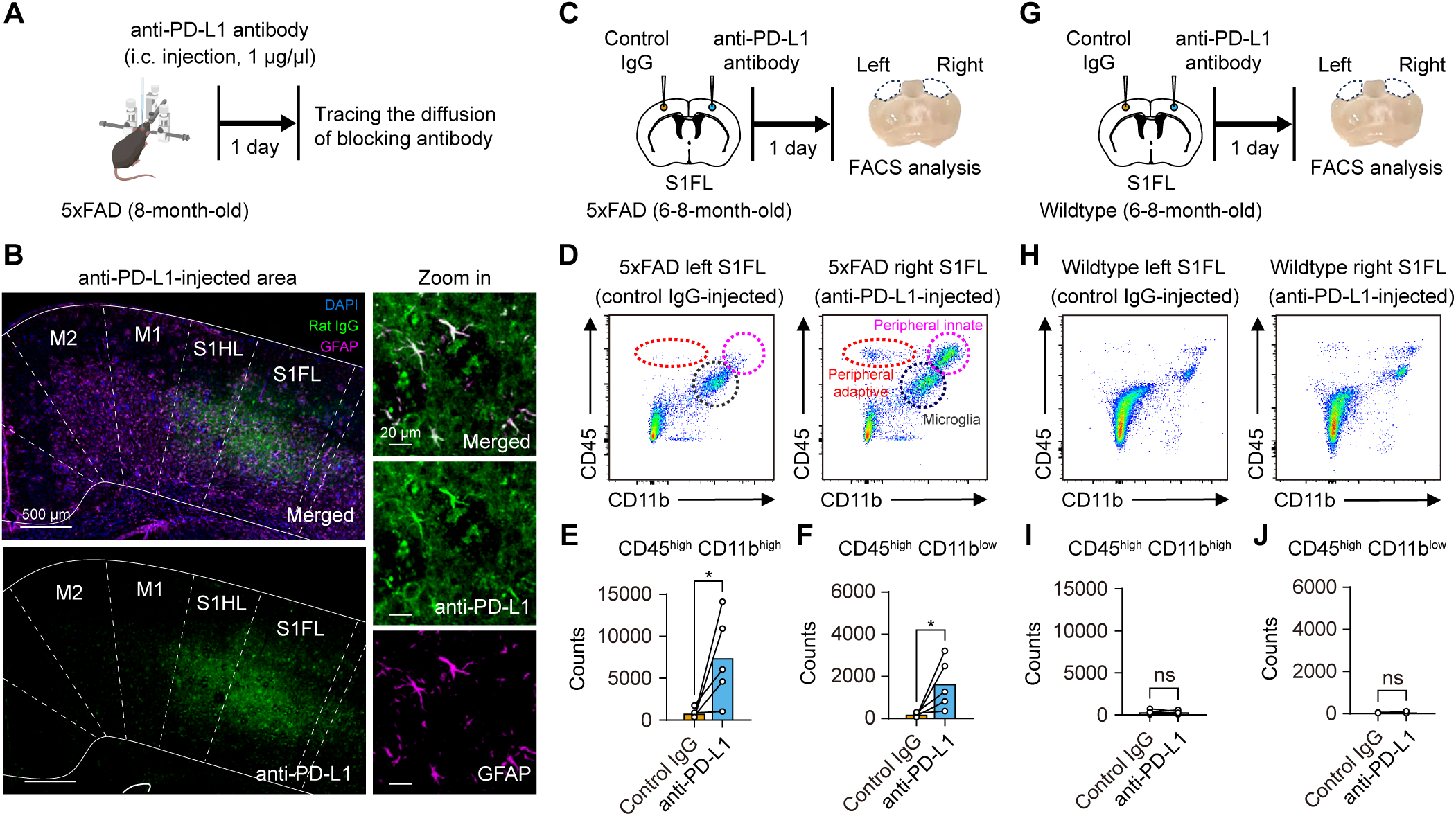
Direct intracortical injection of anti-PD-L1 antibody binds to astrocytes and peripheral immune cells infiltrated in the 5xFAD brain one day post-injection. (**A**) The experimental scheme of the intracortical injection of anti-PD-L1 antibody in 8-month-old 5xFAD mice. (**B**) Representative immunofluorescence images showing the astrocytes and injected materials. Immunostaining for GFAP (magenta) and DAPI (blue), tracking the distribution of the anti-PD-L1 antibody (green) within the somatosensory cortex forelimb region (S1FL; 0 mm AP, ± 2.3 mm ML, − 0.5 mm DV) of 8-month-old 5xFAD mice one day after a single intracortical injection (right, 1 μg/μl). The magnified images (right) show the anti-PD-L1 antibody (green) and GFAP (magenta). (**C**) The experimental scheme of the flow cytometry analysis of the 5xFAD brain tissues by CD11b and CD45 staining in S1FL one day after a single intracortical injection. (**D**) Representative flow cytometry plots showing the peripheral immune cells isolated from the hemispheres of the control IgG-injected area (left hemisphere) and anti-PD-L1-injected area (right hemisphere). (**E**) The quantification of innate immune cell infiltration into the brain (*n* = 5 mice, Wilcoxon signed-rank test; \**p* = 0.043). (**F**) The quantification of adaptive immune cell infiltration into the brain (*n* = 5 mice; Wilcoxon signed-rank test; \**p* = 0.043). (**G**) The experimental scheme of the flow cytometry analysis of the WT brain tissues by CD11b and CD45 staining in S1FL one day after a single intracortical injection. (**H**) Representative flow cytometry plots showing the peripheral immune cells isolated from the hemispheres of the control IgG-injected area (left hemisphere) and anti-PD-L1-injected area (right hemisphere). (**I**) The quantification of innate immune cell infiltration into the brain (*n* = 5 mice; Wilcoxon signed-rank test; p = 0.893, ns, not significant). (**J**) The quantification of adaptive immune cell infiltration into the brain (*n* = 5 mice; Wilcoxon signed-rank test; *p* = 1.000, ns, not significant).

Previous studies have shown that systemic anti-PD-L1 administration through i.p. injection activates the peripheral immune responses in the choroid plexus and recruits the myeloid cells to the CNS (*3*). In contrast, we aimed to investigate the effects of directly modulating the immune checkpoint of the glial cells within the brain. Specifically, the intracortical injection of anti-PD-L1 was performed to determine whether this approach could also promote immune cell infiltration and enhance the brain’s local immune response (Fig. 6, C and G). The flow cytometry analysis of the cells isolated from the injection site of the brain (approximately 1.5 mm³ in volume) one day post-injection revealed a significant increase in peripherally derived immune cells (CD45^high^ and CD11b^low^ lymphocytes and CD45^high^ and CD11b^high^ myeloid cells) in the anti-PD-L1-treated area compared with the control IgG-treated area in the same 5xFAD mouse, indicating substantial immune cell infiltration following the anti-PD-L1 treatment (Fig. 6, D to F). In contrast, the WT mice, which do not express high levels of PD-L1, showed no significant immune cell infiltration following the anti-PD-L1 treatment (Fig. 6, H to J). After seven days post-injection, the earlier increase in infiltrating the immune cells was no longer evident in the anti-PD-L1-treated area, indicating that the infiltration was transient (Fig. S4, A to C). Moreover, no detectable levels of the injected control IgG or anti-PD-L1 antibodies were observed in the brains of 5xFAD mice seven days post-injection (Fig. S4D). These results suggest that the administered anti-PD-L1 antibody has an immediate effect on the local innate cells within the brain tissue but does not affect the infiltrating immune cells. These findings emphasize the critical role of PD-L1 expression and its modulation within the brain in regulating immune cell infiltration and shaping local immune responses in the 5xFAD mouse model.

### Increased chemokine expression and monocyte/macrophage infiltration are induced by the anti-PD-L1 antibody administration into the brain

To determine the immune cell types infiltrating the cortex following anti-PD-L1 treatment, the immune cell populations were profiled using multiparameter flow cytometry, and the effects of anti-PD-L1 on control IgG injections were compared. We first conducted an unbiased uniform manifold approximation and projection (UMAP) analysis to visualize the overall changes in the immune populations (Fig. S5, Fig. S6, Fig. 7A). This analysis revealed a significant increase in immune cells within the anti-PD-L1-treated hemisphere compared with the control IgG-treated hemisphere in the same 5xFAD mouse (Fig. 7, B and C, Fig. S7). Specifically, a proportional analysis revealed a significant increase in Ly6C^+^ monocytes/macrophages (Mo/MΦ) in the anti-PD-L1-treated hemisphere (Fig. 7, D and E). Of note, while the infiltration of peripheral monocytes and macrophages was most evident in the anti-PD-L1-treated hemisphere, we must also consider the possible proliferation or activation of brain-resident perivascular macrophages or other CNS-resident immune cells expressing Ly6C. Although the control IgG-injected hemisphere showed minimal monocyte or macrophage presence (Fig. 6, D to F, Fig. 7B), the observed increase in the Ly6C^+^ cells could comprise both infiltrating peripheral cells and locally proliferating or activated CNS macrophages. Moreover, we observed increases in Ly6C^+^ Mo/MΦ, CD4^+^ conventional T cells (Tconv), and CD4^+^FOXP3^+^ regulatory T (Treg) cells (a non-significant increasing trend was observed), which play critical immunoregulatory roles (Fig. 7F). Further analysis revealed that anti-PD-L1 treatment significantly upregulated chemokines and cytokines, such as CCL5, CXCL10, CCL22, and IL-12p40, which are linked to the enhanced recruitment and infiltration of Mo/MΦ and T cells (Fig. 7G, Fig. S8). We stained the brain sections for the anti-inflammatory cytokine IL-10 and found that it colocalized by infiltrating Ly6C/G^+^ monocytes or macrophages only in the brain area treated with anti-PD-L1 (Fig. 7H). These findings suggest that the local intracortical PD-1/PD-L1 blockade facilitates the recruitment of immunoregulatory cells by enhancing chemokine production, thereby amplifying the immune response and altering the local immune milieu in the brain.

**Figure 7.**
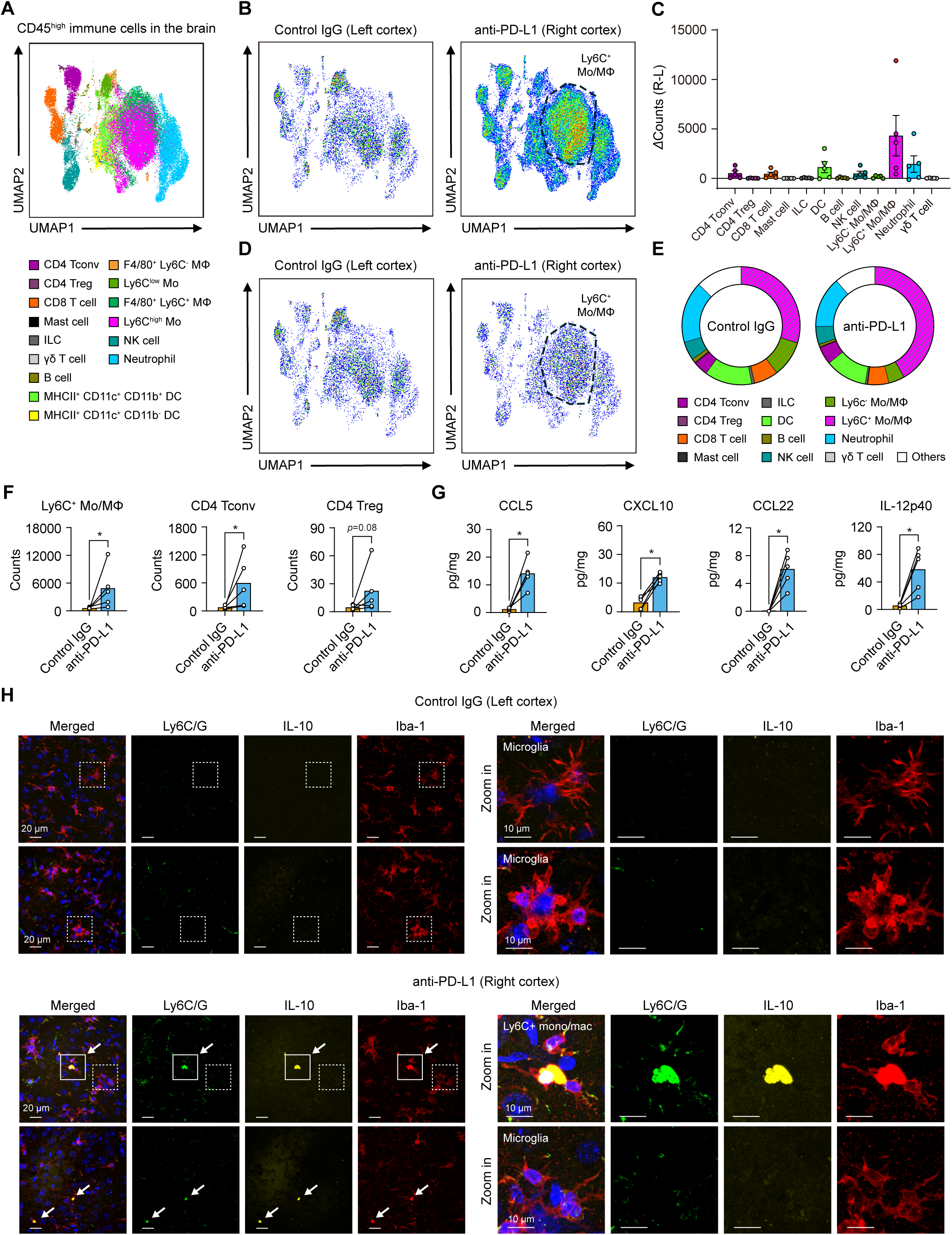
Characterization of the infiltrated immune cell populations and chemokine/cytokine levels in the cortex of 5xFAD mice following the intracortical injection of anti-PD-L1. (**A**) UMAP showing the cell types of the infiltrated immune cells from the cortex of 8-month-old 5xFAD mice injected intracortically with control IgG and anti-PD-L1. (**B**) Cell count plots. (**C**) A graph indicating the number of cells for each cell type, which was obtained by subtracting the number of cells in the control IgG injection group from the number of cells in the anti-PD-L1 injection group (*n* = 5 mice, mean ± s.e.m.). (**D** and **E**) The cell proportion plots from single-cell analysis were down-sampled to 10,000 cells from the original dataset. (**F**) Graphs indicating the number of infiltrated immune cells in the control IgG- and anti-PD-L1-injected cortical regions (*n* = 5 mice, Wilcoxon signed-rank test; Ly6C^+^ Mo/MΦ; \**p* = 0.043, CD4 Tconv; \**p* = 0.043, CD4 Treg; *p* = 0.080, ns, not significant). (**G**) Graphs showing the levels of cytokines and chemokines in the control IgG- and anti-PD-L1-injected cortical regions (*n* = 5 mice, Wilcoxon signed-rank test; CCL5; \**p* = 0.043, CXCL10; \**p* = 0.043, CCL22; \**p* = 0.043, IL-12p40; \**p* = 0.043). (**H**) Representative confocal images showing the colocalization of Ly6C^+^ (green) Iba1^+^ (red) cells and IL-10 (yellow) in the control IgG-injected cortical region (top) and anti-PD-L1-injected cortical region (bottom) of 5xFAD mice (line box; Ly6C/G^+^ monocytes and macrophages, dashed box; microglia, arrow; IL-10^+^ Ly6C/G^+^ monocytes and macrophages). The magnified images (right) showing the microglia or infiltrated immune cells.

## Discussion

This study provides compelling evidence that targeting the PD-1/PD-L1 signaling pathway directly within the brain restores microglial and neuronal functions, providing more insights into the important role of this pathway in the AD brain. Previous studies have shown that systemic anti-PD-L1 administration can reduce Aβ pathology (*2, 3*). However, these investigations did not evaluate its direct functional effects on microglial homeostatic function or neuronal activity. Our results revealed that systemic anti-PD-L1 administration did not promote microglial process convergence (fig. S2), suggesting that peripheral immune activation alone is insufficient to rescue microglial homeostatic function. This may be due to the limited ability of systemically injected antibodies to directly interact with the brain-resident glial cells. Using the intracortical injections of blocking the antibody of the brain intrinsic PD-1/PD-L1 pathway as well as the cell-specific *Pd-l1* knockdown virus, we emphasized the mechanistic interplay between astrocytes, microglia, and neurons, uncovering the role of a critical immune axis in AD.

A key strength of our approach is the use of *in vivo* real-time cellular imaging, which provides direct, visual evidence of the functional improvements in microglial homeostasis, providing more insights beyond the limitations of gene expression analysis alone. Of note, these improvements were observed exclusively following both an intracortical anti-PD-L1 injection and astrocyte-specific *Pd-l1* knockdown, but not with a systemic administration or neuron-specific *Pd-l1* knockdown, emphasizing the importance of targeted brain delivery in promoting glial functional recovery. To achieve the precise cell type-specific knockdown of *Pd-l1* in astrocytes, the pSico system, a Cre-loxP-inducible RNA interference (RNAi) vector, was used. By designing a pSico construct under the control of a GFAP-Cre system, we selectively silenced the *Pd-l1* expression in astrocytes, allowing us to assess its functional role in microglia-astrocyte crosstalk. The knockdown of astrocytic *Pd-l1* replicated the effects of anti-PD-L1 treatment, leading to an enhanced microglial process motility and improved neuronal calcium homeostasis. This specificity emphasizes the astrocyte-driven regulation of PD-L1 signaling in shaping the microglial function. The absence of similar effects in neuron-specific *Pd-l1* knockdown further supports the glial-centric nature of PD-L1 modulation in the pathophysiology of AD. These findings revealed the necessity of precise, cell type-specific manipulations to determine the intricate neuroimmune interactions underlying AD.

The restoration of microglial function led to improvements in neuronal abnormalities, emphasizing the importance of microglia-neuron interactions in AD (*44–46*). Neuronal hyperactivity is a well-documented feature of AD, with numerous studies emphasizing its significance in disease progression (*13–18*). Our intracortical anti-PD-L1 administration effectively addressed the spontaneous neuronal hyperactivity in AD, with its ability to increase the sensory-evoked hemodynamic and neural responses. Moreover, a key distinction to consider is that evoked neural responses represent the collective behavior of neuronal ensembles, rather than the activity of the individual neurons. The decreased hyperactivity of the resting state of the individual neurons may contribute to more coordinated and efficient responses to stimuli at the ensemble level. However, it is important to note that the precise mechanisms linking individual neuronal hyperactivity reduction to improved sensory-evoked neural responses at the population level have not been fully investigated, and further research is needed to fully understand this association. Moreover, restoring microglial function and reducing neuronal hyperactivity may have implications for cognitive outcomes. While systemic PD-1/PD-L1 blockade has been associated with behavioral improvements in AD models (*2, 3*), the cognitive impact of glial PD-1/PD-L1 inhibition needs to be determined. Future research incorporating cognitive assessments could determine whether the observed cellular and molecular changes translate into functional benefits.

Interestingly, peripheral immune cell infiltration was observed only in the 5xFAD brains after anti-PD-L1 administration, suggesting that modulating the PD-1/PD-L1 axis between microglia and astrocytes may also drive this recruitment. While the mechanism by which anti-PD-L1 binding to astrocytes facilitates immune cell entry remains unclear, it is likely that reshaping the astrocyte-microglia interactions leads to the secretion of chemotactic factors (*47, 48*). Further research is needed to confirm the signaling pathways and entry routes involved in this immune cell infiltration. One notable point is that while previous studies have clearly demonstrated peripheral immune cell infiltration following the systemic administration of anti-PD-1 or anti-PD-L1 (*2, 3*), they primarily focused on either phagocytosis or cognitive aspects and did not report microglial functional recovery or enhanced microglia-neuron crosstalk. In contrast, our findings support the idea that beyond peripheral immune cell infiltration, the direct modulation of the PD-1/PD-L1 axis within glial cells influences the functional recovery of both glial cells and neurons in the AD brain.

Additionally, while our study primarily focuses on microglial restoration via intracortical PD-L1 inhibition, it is worth considering whether peripheral monocytes/macrophages also contribute to Aβ clearance. These immune cells are known to exhibit Aβ phagocytic activity (*49, 50*) and express PD-1/PD-L1 (*51, 52*), suggesting that their recruitment into the brain following anti-PD-L1 treatment may play a complementary role in plaque clearance. However, their direct contribution to plaque reduction remains unclear, and we cannot exclude the possibility that anti-PD-L1 acted directly on these infiltrating cells. Nevertheless, our findings demonstrate that astrocyte-specific *Pd-l1* knockdown alone was sufficient to restore microglial homeostasis, indicating that the beneficial effects of anti-PD-L1 are at least partially mediated through astrocyte-microglia interactions rather than solely through infiltrating monocytes. To clarify this further, future studies should assess whether astrocyte-specific *Pd-l1* knockdown using the pSico virus also reduces plaque burden. Such investigations would provide deeper insight into the role of astrocytic PD-L1 signaling in Aβ clearance and help distinguish between its effects on resident versus infiltrating immune cells.

The proper functioning of P2RY12 signaling allows microglia to continuously monitor neuronal activities. The dysfunction of P2RY12 impairs the chemotactic response of microglia to excessive ATP and ADP released from hyperactive neurons, thereby hindering the regulation of neuronal activity (*6*). Our findings suggest that brain-infiltrated monocytes secrete IL-10, which binds to astrocytic IL-10 receptors (IL-10R) to modulate brain inflammation (fig. S9). The increase in the IL-10 levels plays a pivotal role in establishing a more balanced immune environment by reducing the proinflammatory cytokines and neurotoxic factors (*53*). Moreover, IL-10 signaling can stimulate astrocytes to secrete TGF-β, which further amplifies the anti-inflammatory effects and promotes tissue repair (*54, 55*) (Fig. S10). TGF-β, in turn, enhances the expression of P2RY12 on microglia, a receptor critical for their homeostatic and chemotactic functions (*56*). This sequential signaling cascade—from IL-10 secretion by infiltrating monocytes to astrocytic TGF-β release and subsequent P2RY12 upregulation on microglia—provides a coordinated mechanism for restoring the immune balance and neuronal activity in the AD brain. Both systemic anti-PD-L1 treatment (*3*) and our intracortical anti-PD-L1 injection approach led to increased IL-10 levels in the brain, which are correlated with reduced neuroinflammation and improved cognitive function in AD mouse models. These findings emphasize how IL-10-driven TGF-β secretion and subsequent P2RY12 recovery can synergistically repair the disrupted microglia-neuron crosstalk. While our results suggested a protective role for this pathway, other studies have proposed that IL-10 and TGF-β can suppress microglial phagocytosis, potentially impairing the Aβ clearance (*57–59*). This raises the intriguing possibility that the effects of IL-10/TGF-β may be context-dependent, differing between the early and late stages of AD. Rather than a simple pro- or anti-inflammatory role, our data suggest that this signaling axis acts as a dynamic regulator of microglial function, balancing immune modulation and neuroprotection.

Our findings revealed that intracortical anti-PD-L1 administration not only reduces the Aβ plaque size (Fig. 5, A to C) but also increases the plaque-associated microglia (Fig. 5, D and E) and restores homeostatic function in the plaque-associated region, as evidenced by the increased P2RY12 levels (Fig. 3, A to C). Interestingly, in plaques smaller than 150 µm², anti-PD-L1 treatment significantly upregulated P2RY12 in plaque-associated microglia (Fig. S11). This suggests that P2RY12 restoration is not merely due to plaque size reduction but rather driven by an anti-PD-L1-induced increase in astrocytic TGF-β production. Given that Aβ plaques tend to suppress P2RY12 expression (*20, 60*), impair microglial function (*12*), and directly affect neuronal activity (*15*), it is essential to delineate whether these effects are primarily driven by IL-10/TGF-β signaling or occur secondary to enhanced plaque clearance. Furthermore, our results strongly suggest that monocyte-mediated immune modulation, rather than Aβ clearance alone, plays a central role in the restoration of microglial homeostasis following anti-PD-L1 treatment.

A key question is whether the effects of intracortical PD-1/PD-L1 inhibition are transient or sustained. Our data indicated that immune cell infiltration peaks on Day 1 and subsides by Day 7, yet the functional restoration of microglia persists beyond the disappearance of infiltrating monocytes. This timeframe is consistent with previous findings indicating that intracranially administered antibodies tend to exhibit maximal brain retention on Day 1 and largely dissipate on Day 7 (*61*). This suggests that while the administered anti-PD-L1 antibody is cleared within a week, the downstream immune effects persist beyond its presence in the brain. Of note, the sustained impact of PD-L1 inhibition was further supported by our astrocyte-specific *Pd-l1* knockdown experiment using the pSico virus, wherein the continuous suppression of PD-L1 for two weeks led to microglial restoration and functional improvements. This finding reinforces that the observed effects are not solely dependent on the presence of transient antibodies but rather stem from a durable immune reprogramming that persists beyond the initial treatment window. This prolonged effect is likely mediated by IL-10 secreted by recruited monocytes, which triggers an astrocyte-microglia signaling cascade that extends beyond the initial immune response. Although the transient nature of immune infiltration suggests a limited window of direct monocyte influence, the persistence of homeostatic microglial properties suggests a long-lasting shift in the brain’s immune environment.

Another critical insight from our study is that PD-1/PD-L1 blockade restores microglial homeostasis without inducing excessive activation. Unlike the classical immune stimulators that drive proinflammatory responses, PD-L1 inhibition tends to promote microglial process convergence and neuronal surveillance—the hallmarks of homeostatic microglia—without triggering pathological overactivation. This suggests that IL-10 and TGF-β act as key regulators in maintaining the microglial balance, allowing functional restoration, while preventing detrimental hyperactivation. The specificity of this effect on PD-1/PD-L1 signaling inhibition, as opposed to a generalized immune-modulating response, requires a direct comparison with other microglial activators. However, our findings already establish PD-1/PD-L1 blockade as a mechanism that reprograms microglia toward a beneficial, homeostatic state, reinforcing its potential as a strategy for restoring immune balance in AD.

In conclusion, our study establishes a mechanistic association between PD-1/PD-L1 signaling, astrocytic and microglial functions, and neuronal calcium activity in the AD brain. Our approach uniquely combines molecular analysis with direct, functional cellular imaging, providing a more comprehensive and visually intuitive understanding of cellular dynamics in the AD brain. This study provides a basic mechanistic perspective for future research and potential therapeutic interventions, focusing on how targeting brain-innate PD-1/PD-L1 signaling affects functional cellular behaviors in the context of AD pathology.

## Materials and Methods

### Animal models

All experimental procedures were approved by the Institutional Animal Care and Use Committee of Sungkyunkwan University (SKKU). The experiments were conducted in accordance with guidelines established by the Guide for the Care and Use of Laboratory Animals, in compliance with the Animal Protection Law and the Laboratory Animal Act, as regulated by the Korea Animal and Plant Quarantine Agency and the Korea Ministry of Food and Drug Safety. The mouse lines used in the study included the following: C57BL/6J mice, B6.Cg-Tg(APPSwFlLon,PSEN1*M146L*L286V)6799Vas/Mmjax (5xFAD) (*62*), and B6.129P2(Cg)-*Cx3cr1^tm1Litt^*/J (CX3CR1^GFP/GFP^) (*63*) from the Jackson Laboratory. In this study, 4- to 12-month-old 5xFAD mice and age-matched WT littermates were used.

### Intracortical injection

The mice were initially anesthetized using 3% isoflurane for induction and maintained under 1%–1.5% isoflurane throughout the surgery using an isoflurane vaporizer (VetEquip). They were positioned in a stereotaxic frame on a temperature-controlled heating pad (FHC) set at a temperature of 37°C to maintain the body temperature. The head skin of the mouse was sanitized with an alcohol swab (isopropyl alcohol 70%, BD) and then incised using surgical scissors.

A mouse PD-L1-blocking antibody (anti-PD-L1; cat# BE0101, rat IgG2b, κ, clone 10F.9G2, Bio X cell, 1 μg in 1 μL) was administered using a 10 μl Hamilton syringe equipped with a fine glass pipette at a rate of 80 nl/min, connected to a syringe pump (Harvard apparatus). Isotype control IgG antibody (anti-keyhole limpet hemocyanin; cat# BE0090, rat IgG2b, κ, clone LTF-2, Bio X cell, 1 μg in 1 μL) was injected by following the same procedure. The glass pipette remained in place for 5 mins post-injection to minimize backflow. Subsequently, the drilled holes were sealed with bio-compatible surgical elastomer Kwik-Sil (WPI), and the skin was sutured using surgical sutures (B. Braun Surgical); a veterinary adhesive (Vetbond, Fisher Scientific) was applied. The animals were then returned to their home cages. After the injection of the antibody, the mice were allowed to recover with the application of antibiotics (Baytril; 5 mg/kg, s.c. injection) and analgesics (Meloxicam; 1 mg/kg, s.c. injection) and were closely monitored for any signs of distress.

For the *in vivo* two-photon experiments, injections were administered into the right primary somatosensory cortex of the forelimb region (S1FL) through a small craniotomy performed at a 40-degree angle. This approach was chosen to avoid major blood vessels and minimize potential tissue damage above the injection site during the acute craniotomy performed for imaging. For immunostaining and immunoassays, small craniotomies were performed over the S1FL region in both hemispheres. The left hemisphere was injected with a control isotype antibody, whereas the right hemisphere received an anti-PD-L1 antibody. To assess the immune cell populations, five injection sites were targeted in each hemisphere, totaling 10 sites, to ensure sufficient cell numbers for FACS sorting.

### Virus for cell-specific *Pd-l1* gene silencing

For astrocyte-specific *Cd274* gene silencing, a 1:1 mixture of AAV5-GFAP-mCherry-Cre (titer: 2.49 × 10^12^ GC/ml, IBS virus facility) and AAV5-pSico-*Cd274*-BFP (titer: 2.28 × 10^12^ GC/ml, IBS virus facility) was used. As a control, AAV-pSico-scrambled shRNA-BFP (titer: 2.22 × 10^12^ GC/ml, IBS virus facility) was mixed instead of AAV-pSico-Cd274-BFP. In the case of excitatory neuron-specific *Cd274* gene silencing, AAV9-CaMK2a-Cre (titer: 6.35 × 10^12^ GC/ml, IBS virus facility) and AAV9-pSico-*Cd274*-mCherry (titer: 2.44 × 10^13^ GC/ml, IBS virus facility) were diluted in 0.1 M phosphate-buffered saline (PBS) to adjust approximately 3 × 10^12^ GC/ml. Moreover, the diluted viruses were then mixed in equal volumes and used for injection. As a control, AAV-pSico-scrambled shRNA-mCherry (titer: 7.64 × 10^13^ GC/ml, IBS virus facility) was diluted in PBS and mixed instead of AAV-pSico-*Cd274*-mCherry.

### Cranial window surgery in mice

Initially, the mice were anesthetized using 3% isoflurane for induction and maintained under 1%–1.5% isoflurane throughout the surgery. The body temperature was carefully maintained at 37°C using a temperature-controlled heating pad (FHC). A cranial window was surgically created over the cortex to facilitate imaging, with the dura mater being left intact. The exposed cortex was hydrated with HEPES-buffered saline (135 mM NaCl, 5 mM KCl, 10 mM HEPES, 10 mM glucose, 2 mM CaCl_2_, 2mM MgSO_4_) or HEPES-buffered saline-soaked GelFoam sponges (MS0005, Ethicon). After confirming the absence of microbleeding, a glass coverslip was placed over the exposed cortex (4-mm diameter, Deckglaser). To stabilize the cranial window and prevent animal head movements during imaging, a custom-made metal holding frame was affixed to the skull. This frame also allowed for the adjustment of the cranial window angle to ensure that it remained perpendicular to the microscope objective axis. Throughout the experiments, the physiological parameters, such as heart rate, SpO_2_, and respiration rate, were continuously monitored using a paw sensor (PhysioSuite, Kent Scientific) to ensure they remained within normal ranges. At the conclusion of all the experimental procedures, the mice were euthanized by CO_2_ inhalation in a closed chamber.

### *In vivo* two-photon imaging

*In vivo* two-photon imaging was performed to visualize the microglial dynamics and spontaneous neuronal activity in live mice. A two-photon microscope (TCS SP8 MP, Leica Microsystems) equipped with a Ti:sapphire laser (Chameleon Vision II, Coherent) was used for brain imaging. Moreover, imaging was performed using a 25× water-immersion objective lens (NA 0.95, Leica Microsystems) and a green bandpass emission filter set at 520 ± 50 nm. Isoflurane anesthesia can affect the homeostatic function of microglia and neuronal activity (*64*). To minimize these effects, the mice were kept under light isoflurane anesthesia, with the concentration carefully maintained at a low level (∼0.8%) throughout the imaging process, as in previous studies (*15, 17, 18, 65, 66*).

For the *in vivo* two-photon imaging of microglia, 5xFAD mice were crossbred with CX3CR1-GFP mice, wherein the microglia were genetically modified to express GFP under the control of the CX3CR1 promoter. This crossbreeding produced offspring with fluorescence specifically in microglia, enabling targeted imaging and the analysis of microglial dynamics within the 5xFAD model. Imaging was conducted at wavelengths tuned to 920 nm, focusing on the regions deeper than 250 μm in the cortex, which had been injected with either anti-PD-L1 or isotype control antibodies. For laser-induced injury, the laser was focused at ∼50 mW for 60 s. The microglial responses were calculated using the following formula: (If (t) − If (0)) / Is (t), wherein If represents the intensity at the focus (with a diameter of 50 μm), whereas Is represents the intensity of the surrounding area, forming a donut-like shape due to the subtraction of the focus (*67*).

For the *in vivo* two-photon imaging of the neurons, the pENN.AAV.CamKII.GCaMP6f. WPRE.SV40 virus (Plasmid #100834, AAV9 (titer: 3.1 x 10^13^ GC/ml)) was co-delivered with either anti-PD-L1 or isotype control antibodies to visualize excitatory neuronal calcium activity. Spontaneous Ca^2+^ fluorescence signals from the cortical layer I/III were recorded at a frequency of ∼15 Hz. Imaging was conducted across multiple regions per mouse, with each region recorded for at least 2 min. The cell region of interests (ROI) were drawn around individual neuronal somata, and then relative GCaMP6f fluorescence changes over time were generated for each ROI. The neurons were classified based on their activity rates, with hyperactive neurons defined as those exhibiting ≥4 transients per minute (*15*).

### IOS imaging

Hemodynamic responses in the right S1FL region were recorded using an optical imaging system (Imager 3001-Celox, Optical Imaging Ltd.) following forepaw electrical stimulation. To ensure a stable imaging environment, the exposed cortical surface was enclosed by a dental resin wall and filled with silicone oil (Sigma). The region was illuminated using an LED lamp (CLS150, Leica Microsystems), while the images were captured at 10 Hz using a fast-acquisition camera (Photonfocus AG) with 50 mm tandem lenses. To measure the total hemoglobin (tHb), reflected light was filtered at 546 ± 30 nm, the isosbestic point for oxyhemoglobin and deoxyhemoglobin absorption. The field of view had a pixel resolution of 372 × 372 pixels, with a pixel size of 8 μm. For electrical stimulation, 500 μs pulses at 4 Hz and 0.5 mA were applied to the left forepaw for 20 s using a pulse stimulator (Master-9, A.M.P.I.) and a current generator (ISO-Flex, A.M.P.I.). The hemodynamic intensity changes were calculated per trial, with the baseline defined as the 5 s preceding the stimulation onset. A 21 × 21 pixels ROI was selected for analysis. The time-series data were temporally averaged at each time point, and the peak amplitude was determined within the stimulation period. The onset time was defined as the point where the intensity changes exceeded the mean baseline plus 2 standard deviations.

### LFP recording

IOS imaging was used to determine the activated area within the right S1FL region, guiding the insertion site for electrophysiological recordings. A low-impedance electrode (∼0.5 MΩ, FHC, Inc.) was positioned at the center of the activated region, targeting layer II/III of the somatosensory cortex at a depth of 300 μm. To minimize electrical noise and external interference, a grounding screw was placed in the left olfactory bulb region and connected to the ground wire, ensuring stable and reliable LFP recordings. Electrophysiological activity was recorded at 40 kHz using a Plexon system (Plexon Inc.). For the LFP analysis, the raw signals were bandpass-filtered between 0.5 and 200 Hz with 60-Hz notch filtering (55–65 Hz) to remove the electrical noise. LFP data processing and spectral analysis were performed using the Chronux toolbox, an open-source platform for time-series neural data analysis.

### Multiparameter flow cytometry analysis and cell sorting

The mice were deeply anesthetized with 2.5% avertin (400 mg/kg, i.p., cat# 240486, Sigma) and perfused with PBS. The brains were then extracted, mechanically minced, and incubated at a temperature of 37°C for 30 min in a dissociation buffer containing 0.1 mg/mL DNase I (Roche), 1 mg/mL collagenase type IV (Gibco), and 2% FBS in DMEM Medium. The digested brain tissue was gently mashed through a 70-μm strainer and centrifuged at 750 g for 5 min at a temperature of 4°C in a swinging bucket rotor. Pellets were resuspended with 35% Percoll in PBS (Cytiva) and centrifuged at 1120 g for 20 min at room temperature (RT) to remove myelin debris. The pellets were resuspended in DMEM with 10% FBS for further analysis.

For the flow cytometry analysis, the isolated cells were surface-stained with the following antibodies: CD45-BUV395 (1:200 dilution, cat# 564279, clone 30-F11, BD), CD8-BUV805 (1:200 dilution, cat# 612898, clone 53–6.7, BD), CD62L-BUV737 (1:200 dilution, cat# 612833, clone MEL-14, BD), CD44-BUV615 (1:200 dilution, cat# 751414, clone IM7, BD), F4/80-BUV661 (1:200 dilution, cat# 750643, clone T45–2342, BD), MHCII-BUV496 (1:200 dilution, cat# 750281, clone M5/114.15.2, BD), NK1.1-BUV563 (1:200 dilution, cat# 741233, clone PK136, BD), TCRγδ-BV605 (1:200 dilution, cat# 107502, clone UC7– 13D5, BioLegend), TCRβ-BV650 (1:200 dilution, cat# 109207, clone H57–597, BioLegend), Ly6C-BV785 (1:200 dilution, cat# 128041, clone HK1.4, BioLegend), CD127-PE-Cy5 (1:200 dilution, cat# 135015, clone A7R34, BioLegend), CD11b-PerCP-Cy5.5 (1:200 dilution, cat# 101227, clone M1/70, BioLegend), CD11c-PE-Cy7 (1:200 dilution, cat# 117317, clone N418, BioLegend), CD19-APC-R700 (1:200 dilution, cat# 565473, clone 1D3, BD), CD4-APC-Cy7 (1:200 dilution, cat# 100413, clone GK1.5, BioLegend), C-kit-PerCP (1:200 dilution, cat# 105801, clone 2B8, BioLegend), and FcεRIα-AF647 (1:200 dilution, cat# 134309, clone MAR-1, BioLegend). Moreover, LIVE/DEAD™ Fixable Aqua (Thermo Fisher Scientific, USA) was added to a staining solution to discriminate the dead cells. After incubation at a temperature of 4 °C for 30 min, the cells were fixed and permeabilized with eBioscience™ Foxp3/Transcription Factor Staining Buffer (Invitrogen, USA) and were stained with the following antibodies at RT for 40 min: FOXP3-AF488 (1:100 dilution, cat# 53–5773–82, clone FJK-16s, Invitrogen), HELIOS-PB (1:100 dilution, cat# 137210, clone 22F6, BioLegend), and RORγt-PE (1:100 dilution, cat# 12–6981–82, clone B2D, Invitrogen). After staining, the cells were washed with PBS, and data acquisition was performed using a Sony ID7000 (Sony).

For cell sorting, the isolated cells were stained with the following antibodies: CD11b-FITC (1:200 dilution, cat# 101205, clone M1/70, BioLegend), ACSA-2-PE (1:200 dilution, cat# 130–123–284, clone IH3–18A3, Miltenyi Biotec), and CD45-APC-Cy7 (1:200 dilution, cat# 103115, clone 30-F11, BioLegend). After staining, the cells were washed with PBS, and the cell pellets were resuspended in DMEM with 10% FBS. Moreover, DAPI was added to distinguish the dead cells. Microglia and astrocytes were sorted on the Sony MA900 cell sorter (Sony).

Extended Data Figure 3 and Extended Data Figure 4 show the gating strategies for the flow cytometric analysis and cell sorting. The flow cytometric data were analyzed in FlowJo v10.9.0 (BD). The UMAP algorithm for dimensionality reduction was applied using the UMAP plugin (v3.1) available on FlowJo.

### RNA isolation and reverse transcription-quantitative PCR analysis

For the quantification of mRNA, mRNA was extracted from the sorted cells using the RNeasy Micro Kit (Qiagen), while PrimeScript™ II 1st strand cDNA Synthesis Kit (Takara) was used for reverse transcription. PowerUp™ SYBR® Green PCR Master Mix (Thermo Fisher) and QuantStudio3 (Applied Biosystems) were used for quantitative PCR analysis with the following primers targeting Tgfb1: F: 5′-CTCCCGTGGCTTCTAGTGC-3′ and R: 5′-GCCTTAGTTTGGACAGGATCTG-3′.

### Multiplex chemokine and cytokine analysis

For multiplex chemokine and cytokine analysis, the anti-PD-L1- and control IgG-injected cortical regions were dissected, and the brain lysate was prepared with cell lysis buffer (Cell signaling) by homogenization and sonication. Chemokine and cytokine in plasma were measured using LEGENDplex™ Mouse Proinflammatory Chemokine Panel (cat# 740007, BioLegend) for CCL5,CCL20, CCL11, CCL17, CXCL1, CCL2, CXCL9, CXCL10, CCL3, CCL4, CXCL13, CXCL5, and CCL22; Mouse Proinflammatory Chemokine Panel 2 (cat# 741068, BioLegend) for CCL24, CCL8, CXCL11, CX3CL1, CCL7, XCL1, CXCL2, and CCL9; Mouse Th Cytokine Panel (cat# 741044, BioLegend) for IFN-γ, IL-5, TNF-α, IL-2, IL-6, IL-4, IL-10, IL-9, IL-17A, IL17F, IL-22, and IL-13; and Mouse Cytokine Panel 2 (cat# 740134, BioLegend) for IL-1α, IL-1β, IL-3, IL-7, IL-11, IL-12p40, IL-12p70, IL-23, IL-27, IL-33, IFN-β, GM-CSF, and TSLP. For each sample, more than 300 beads were recorded using BD FACSCelesta (BD) and analyzed using the Data Analysis Software Suite for LEGENDplex, a free cloud-based program (https://legendplex.qognit.com/).

### Immunostaining and histological analysis

The mice were anesthetized with 2.5% avertin (400 mg/kg, i.p.) and perfused with 0.1 M PBS followed by 4% paraformaldehyde (PFA) using a peristaltic pump (P-1500, Harvard apparatus) after receiving an intracortical injection on either Day 1 or Day 7. The perfused brains were placed in 4% PFA for 24 h at a temperature of 4 °C, followed by immersion in 30% sucrose in PBS for three days. Forty µm-thick coronal brain tissue sections were permeabilized with 0.5% Triton X-100 in PBS for 10 min at RT, blocked with 10% donkey serum, and incubated with the primary antibody overnight at a temperature of 4 °C. The following primary antibodies were used: rabbit anti-PD-1 (1:300 dilution, cat# PRS4065, Sigma), rat anti-PD-L1 (1:500 dilution, cat# BE0101, Bio X Cell), rabbit anti-Iba1 (1:300 dilution, cat# 234–004, Wako), goat anti-Iba1 (1:300 dilution, cat# ab5076, Abcam), mouse anti-GFAP (1:400 dilution, cat# MAB360, Millipore), goat anti-GFAP (1:400 dilution, cat# ab53554, Abcam), mouse anti-NeuN (1:800 dilution, cat# ab104224, Abcam), rabbit anti-NeuN (1:800 dilution, cat# 234009, Abcam), rat anti-IL-10 (1:100 dilution, cat# ab189392, Abcam), rat anti-CD210 (IL-10 receptor) (1:500 dilution, cat# 112706, BioLegend), rabbit anti-P2RY12 (1:1000 dilution, cat# ab300140, Abcam), rabbit anti-GFP (1:800 dilution, cat# A-11122, Invitrogen), mouse anti-β-amyloid 1–16 (6E10) (1:1000 dilution, cat# 803014, BioLegend), and rat anti-CD68 (1:200 dilution, cat# MCA1957, Bio-Rad). The following day, the sections were washed in PBS and then incubated in secondary antibodies for 2 h at RT. The following secondary antibodies were applied respectively: anti-rat IgG Alexa488 (donkey, 1:400 dilution, cat# A21208, donkey, Invitrogen), anti-goat IgG Alexa555 (donkey, 1:400 dilution, cat# A21432, Invitrogen), anti-goat IgG Alexa633 (donkey, 1:400 dilution, cat# A21082, Invitrogen), anti-mouse IgG Alexa488 (donkey, 1:400 dilution, cat# A21202, Invitrogen), anti-mouse IgG Alexa568 (donkey, 1:400 dilution, cat# A10037, Invitrogen), anti-rabbit IgG Alexa488 (donkey, 1:400 dilution, cat# A21206, Invitrogen), anti-rabbit IgG Alexa568 (donkey, 1:400 dilution, cat# A10042, Invitrogen), and anti-rabbit IgG Alexa647 (donkey, 1:400 dilution, cat# A31573, Invitrogen). For Thioflavin S staining (ThioS, diluted in 50% ethanol, cat# T1892, Sigma), the brain sections were incubated in 0.5% ThioS for 10 min at RT. The monocytes were stained via the tail vein injection of an Alexa Fluor 488-conjugated Ly6C/G (Gr-1) antibody (rat, 0.12 mg/kg, cat# 53–5931–82, eBioscience). Nuclear counterstaining was performed with 100 ng/ml DAPI solution (1:10,000 dilution, cat# D9542, Sigma) in PBS for 10 min.

Confocal microscopy (TCS SP8, Leica Microsystems) was used for fluorescence imaging, and the quantitative analysis of the staining intensity and cellular localization was conducted using the Fiji67 and IMARIS (Bitplane) software. To analyze the cell-specific expression of PD-1 and PD-L1, the fluorescence images obtained using the anti-Iba1, anti-GFAP, and anti-NeuN primary antibodies were converted to binary images, and the immunoreactivity of PD-1 and PD-L1 within the ROIs was calculated. The assessment of microglial phagocytosis and Aβ clearance was conducted with the colocalization of the microglia-specific lysosome marker (CD68) and Aβ (6E10) on the confocal optical sections using IMARIS with its module (*68*). The brightest approximately 2% of the pixels in each channel were evaluated to determine the threshold for colocalization in an unbiased manner, eliminating any problems caused by inevitable imaging variations.

### Statistical analysis

The statistical analyses were conducted using GraphPad Prism 9.5.1 (GraphPad Software, USA) or IBM SPSS Statistics 19 (SPSS Inc., USA). The two groups were compared using the Mann–Whitney U test, with the Wilcoxon signed-rank tests used for paired data. For comparisons involving more than three groups, Dunn’s multiple comparison test was used, with the Bonferroni post-hoc procedure applied for pairwise comparisons. The relationship between the two variables was assessed using Spearman’s correlation coefficient. Ordinary least squares regression was used to fit a linear model to the data points. Furthermore, statistical significance is indicated in the figures by * for p < 0.05, ** for p < 0.01, *** for p < 0.001, and **** for p < 0.0001.

## Acknowledgments

We would like to thank Hyouna Yoo, Euna Lee, Jae Hoon Jeong, Hyojung Kim, and Bok-Man Kang of IMNEWRUN Inc. for their helpful discussions and comments on the manuscript. We also thank Gyeong Eon Kim for proofreading the manuscript.

## Author contributions

Conceptualization: TP, HJK, and MS

Project administration: TP, MS

Stereotaxic injections: TP, SWC, SL

*In vivo* two-photon imaging and data analysis: TP

IOS imaging and data analysis: TP

LFP recording and data analysis: TP and SB

Immunofluorescence staining: TP and SB

Cellular and molecular immunoassays: LC and HKK

Supervision: YHK, JL, HJK, HKK, and MS

Writing—original draft: TP, LC, HKK, and MS

Writing—review & editing: TP, LC, SL, SWC, SB, CC, JL, HJK, HKK, and MS

## Data and materials availability

All data needed to evaluate the conclusions in the paper are present in the paper and/or the Supplementary Materials.

---

**Fig. S1.**
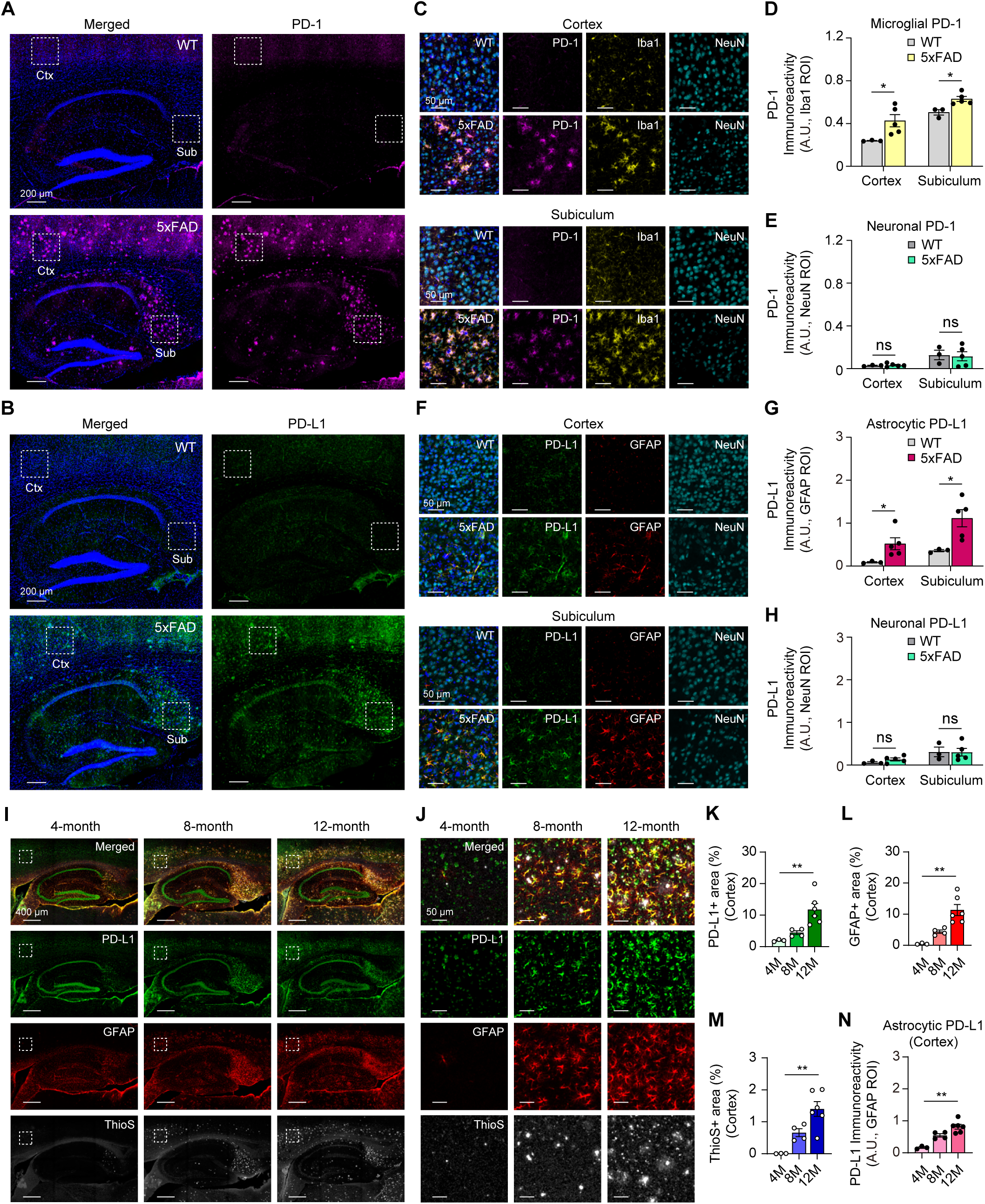
Upregulated expression of microglial PD-1 and astrocytic PD-L1 was found in 8-month-old 5xFAD mice. (**A**) Representative confocal microscopy images of PD-1 immunoreactivity (magenta) stained with DAPI (blue) in the control brains and 5xFAD mouse brains. Ctx indicates the cortex. Sub indicates the subiculum. (**B**) Representative confocal microscopy images of PD-L1 immunoreactivity (green) stained with DAPI (blue) in the control brains and 5xFAD mouse brains. (**C**) The magnification images of the white dashed box in (A). Colocalization is shown in the merged images. Brain slices were stained with DAPI (blue), PD-1 (magenta), the microglial marker Iba1 (yellow), and the neuronal marker NeuN (cyan). (**D**) Quantification indicating the microglial PD-1 expression in the 5xFAD brain (WT; *n* = 3 mice, 5xFAD; *n* = 5 mice, Mann–Whitney U test; cortex; \**p* = 0.025, subiculum; \**p* = 0.025, mean ± s.e.m.). (**E**) Quantification indicating the neuronal PD-1 expression in the 5xFAD brain (Mann–Whitney U test; cortex; *p* = 0.881, subiculum; *p* = 0.655, mean ± s.e.m., ns, not significant). (**F**) The magnification images of the white dashed box in (B). Colocalization is shown in the merged image. Brain slices were stained with DAPI (blue), PD-L1 (green), astrocytic marker GFAP (red), and NeuN (cyan). (**G**) Quantification indicating the astrocytic PD-L1 expression in the 5xFAD brain (Mann–Whitney U test; cortex; \**p* = 0.025, subiculum; \**p* = 0.025, mean ± s.e.m.). (**H**) Quantification indicating the neuronal PD-L1 expression in the 5xFAD brain (Mann–Whitney U test; cortex; *p* = 0.297, subiculum; *p* = 0.881, mean ± s.e.m., ns, not significant). (**I**) Representative images showing the age-dependent changes in PD-L1 (green), GFAP (red), and Thioflavin S (ThioS) (gray) in the 5xFAD mouse brains at different disease stages. (**J**) The magnification images of the white dashed box in (I) (cortex). (**K**-**M**) The quantification of the PD-L1, GFAP, and Aβ (ThioS) expression levels across time points (4M = 4-month-old; *n* = 3 mice, 8M = 8-month-old; *n* = 4 mice, 12M = 12-month-old; *n* = 6 mice, Kruskal-Wallis test with Dunn’s post-hoc comparisons test; PD-L1 4M vs 8M; *p* = 0.7179, PD-L1 4M vs 12M; \*\**p* = 0.0061, PD-L1 8M vs 12M; *p* = 0.1401; GFAP 4M vs 8M; *p* = 0.7179, GFAP 4M vs 12M; \*\**p* = 0.0061, GFAP 8M vs 12M; *p* = 0.1401; ThioS 4M vs 8M; *p* = 0.5361, ThioS 4M vs 12M; \*\**p* = 0.0091, ThioS 8M vs 12M; *p* = 0.2923, mean ± s.e.m.). (**N**) Quantification indicating the age-dependent astrocytic PD-L1 expression in the 5xFAD brain (Kruskal-Wallis test with Dunn’s post-hoc comparisons test; 4M vs 8M; *p* = 0.5361, 4M vs 12M; \*\**p* = 0.0091, 8M vs 12M; *p* = 0.2923, mean ± s.e.m.).

**Fig. S2.**
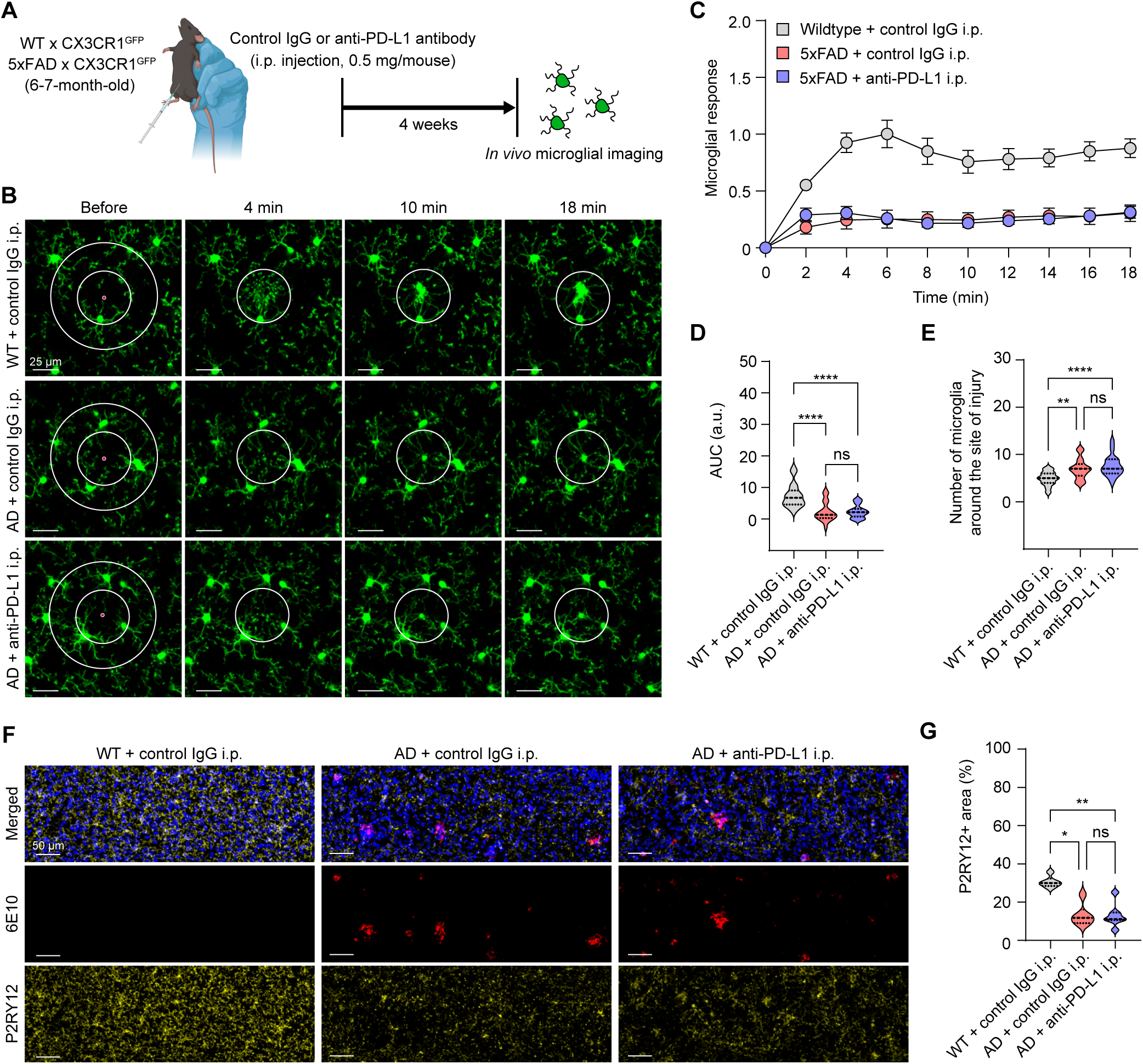
Systemic injection of anti-PD-L1 does not alter the microglial process convergence in response to local brain injury in 5xFAD mice. (**A**) The experimental scheme of *in vivo* two-photon (2P) microglial time-lapse imaging with focal laser injury four weeks after a single i.p. injection of antibody in the WT/CX3CR1^GFP^ or 5xFAD/CX-3CR1^GFP^ mice. (**B**) Representative time-lapse imaging of laser injury-targeted microglial process convergence. (**C**) Time course of microglial process kinetics toward a laser-induced lesion (WT + control IgG i.p.; *n* = 26, 5xFAD + control IgG i.p.; *n* = 25, 5xFAD + anti-PD-L1 i.p.; *n* = 24). Microglial responses = If(t)-If(0)/Is(t); If indicates the intensity of the focus, whereas Is indicates the intensity of the surrounding area (mean ± s.e.m.). (**D**) The AUC of the accumulated graphs in (C) (Kruskal-Wallis test with Dunn’s post-hoc comparisons test; WT + control IgG i.p. vs. AD + control IgG i.p.; \*\*\*\**p* < 0.0001, WT + control IgG i.p. vs. AD + anti-PD-L1 i.p.; \*\*\*\**p* < 0.0001, AD + control IgG i.p. vs. AD + anti-PD-L1 i.p.; *p* = 1.000, ns, not significant). (**E**) Number of microglia around the site of focal injury within 100 μm (Kruskal-Wallis test with Dunn’s post-hoc comparisons test; WT + control IgG i.p. vs. AD + control IgG i.p.; \*\**p* = 0.0019, WT + control IgG i.p. vs. AD + anti-PD-L1 i.p.; \*\*\*\**p* < 0.0001, AD + control IgG i.p. vs. AD + anti-PD-L1 i.p.; *p* = 0.9349, ns, not significant). (**F**) Representative images of P2RY12 (yellow) and 6E10 (red) four weeks after a single i.p. injection of the antibody. (**G**) The quantitation of the P2RY12-positive area (WT + control IgG i.p.; *n* = 6 mice, 5xFAD + control IgG i.p.; *n* = 6 mice, 5xFAD + anti-PD-L1 i.p.; *n* = 7 mice, Kruskal-Wallis test with Dunn’s post-hoc comparisons test; WT + control IgG i.p. vs. AD + control IgG i.p.; \**p* = 0.0104, WT + control IgG i.p. vs. AD + anti-PD-L1 i.p.; \*\**p* = 0.0072, AD + control IgG i.p. vs. AD + anti-PD-L1 i.p.; *p* = 1.000, ns, not significant).

**Fig. S3.**
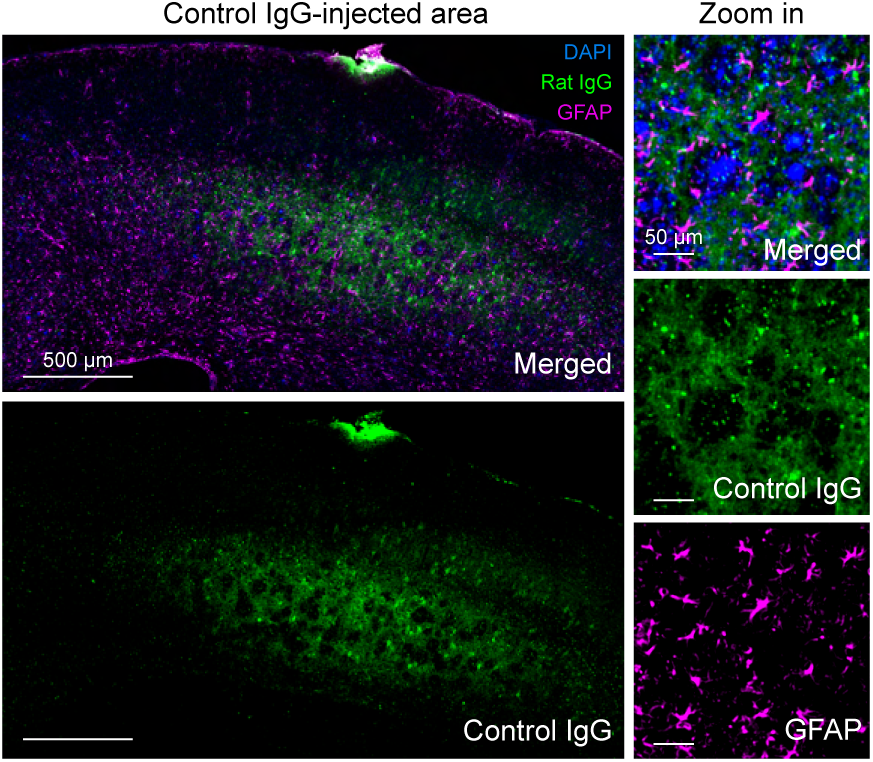
Distribution of the injected control IgG in the brains of 5xFAD mice. Representative immunofluorescence images showing the astrocytes and injected materials. Immunostaining for GFAP (magenta) and DAPI (blue), tracking the distribution of the control IgG (green) within the S1FL of 8-month-old 5xFAD mice one day after a single intracortical injection (left, 1 μg/μl). The magnified images (right), indicating the DAPI (blue), control IgG (green), and GFAP (magenta).

**Fig. S4.**
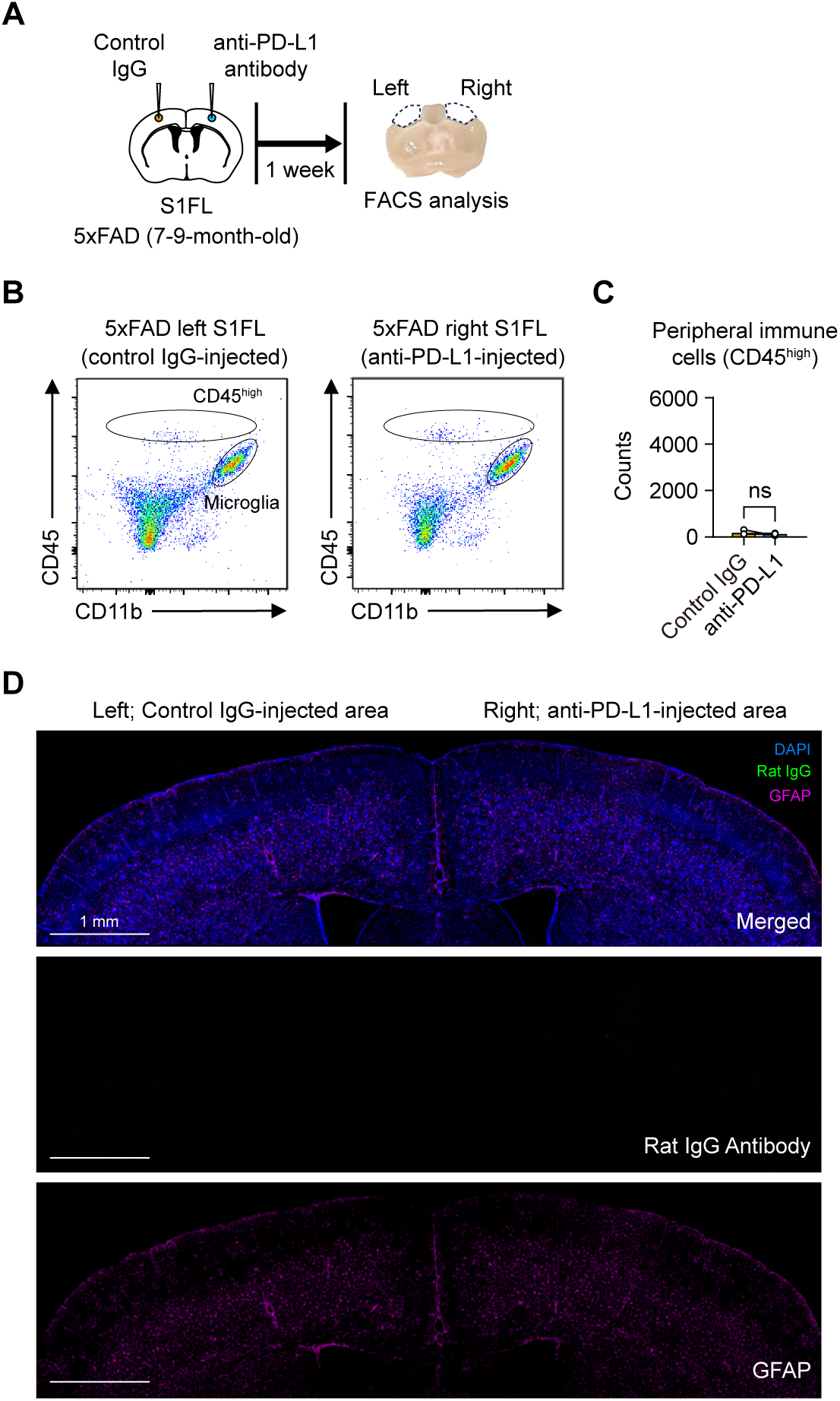
Resolution of early immune cell infiltration and absence of injected antibody in the brain of 5xFAD mice by seven days post-injection. (**A**) The experimental scheme of flow cytometry analysis of 5xFAD brain tissues by CD11b and CD45 staining in S1FL at seven days after single intracortical injection. (**B**) Representative flow cytometry plots showing peripheral immune cells isolated from the hemispheres of control IgG-injected area (left hemisphere) and anti-PD-L1-injected area (right hemisphere). (**C**) The quantification of immune cell infiltration into the brain (*n* = 5 mice, Wilcoxon signed-rank test; *p* = 0.225, ns, not significant). (**D**) Representative immunofluorescence images showing astrocytes and injected materials. Immunostaining for GFAP tracking the distribution of the control IgG (left hemisphere) and anti-PD-L1 (right hemisphere) within the S1FL of 8-month-old 5xFAD mice, seven days after a single intracortical injection (1 μg/μl, DAPI; blue, rat IgG; green, GFAP; magenta).

**Fig. S5.**
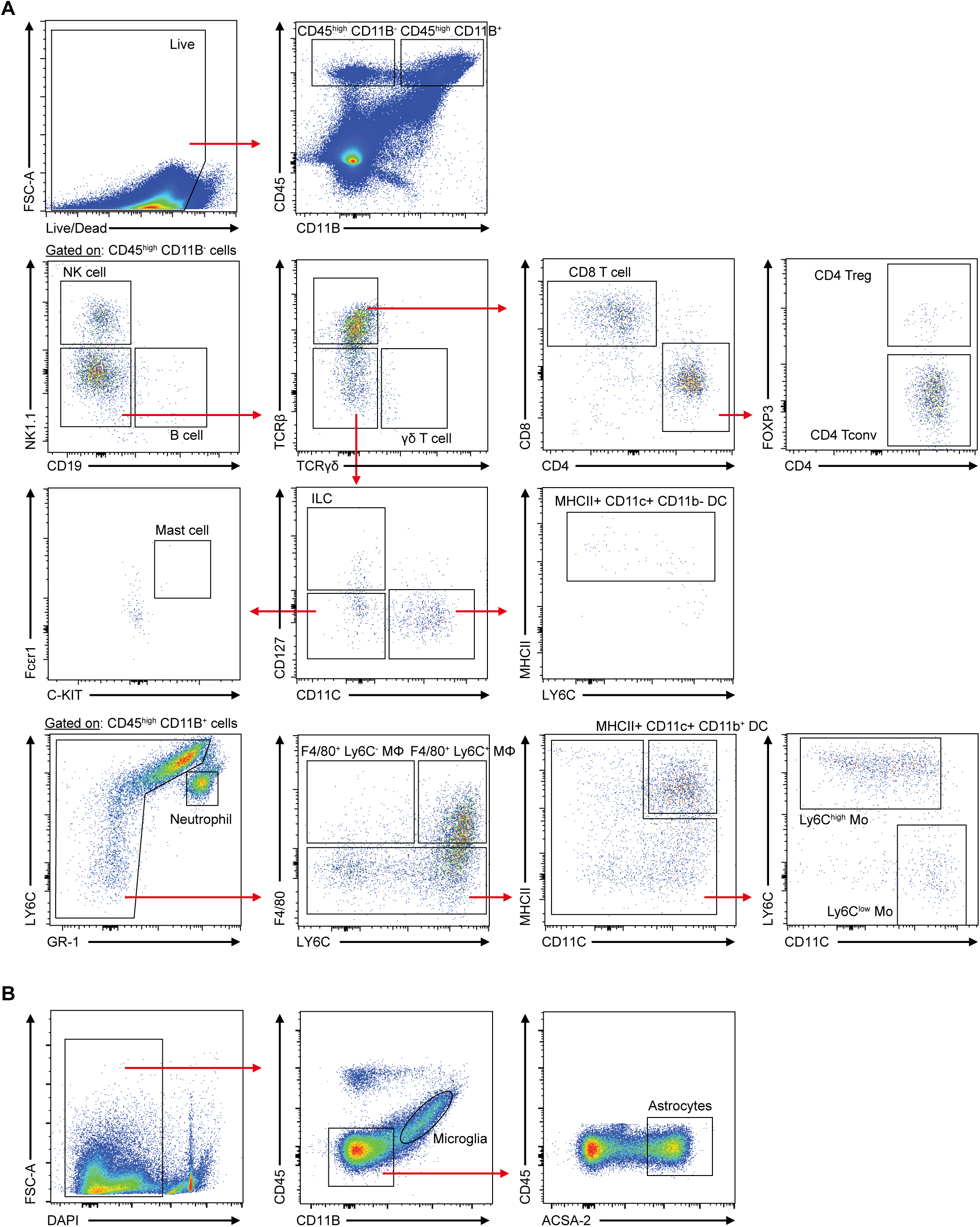
Flow cytometry gating strategy used to define the cell populations from the mouse brain. (**A**) Gating strategy for the flow cytometry analysis of the brain-infiltrated immune cells after anti-PD-L1 injection. (**B**) Gating strategy for microglia and astrocyte cell sorting.

**Fig. S6.**
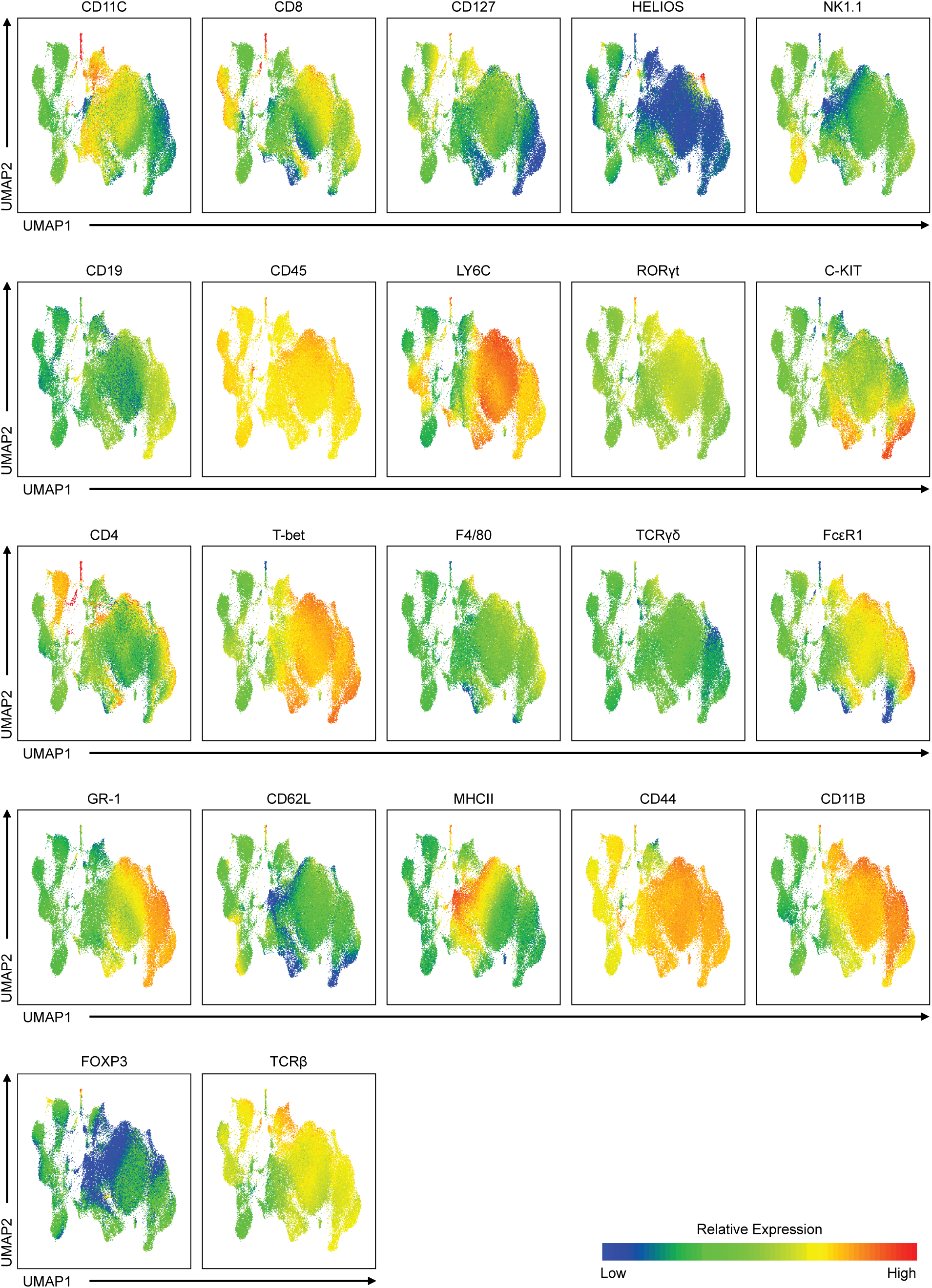
UMAP plot showing the expression levels of the cell type-specific markers used for flow cytometry analysis. CD11C, expressed on dendritic cells. CD8, found on cytotoxic T cells. CD127, IL-7 receptor, identifying memory and naïve T cells. HELIOS, a transcription factor linked to Treg cells. NK1.1, a marker for natural killer (NK) cells. CD19, B cell-specific marker. CD45, a pan-leukocyte marker, indicative of all hematopoietic cells. LY6C is a marker for infiltrating monocytes, which can differentiate into macrophages that assist in tissue repair and debris clearance. RORγt, a transcription factor specific to Th17 cells. C-KIT, a marker of hematopoietic stem cells and progenitor cells. CD4, expressed on helper T cells and aids in coordinating immune responses. T-bet, a transcription factor associated with Th1 cells and cytotoxic T cells. F4/80, a marker for identifying macrophages. TCRγδ, a marker for gamma-delta T cells. FcεRI, a high-affinity IgE receptor, indicating mast cells and basophils. GR-1, a granulocyte marker, typically expressed on neutrophils and monocytes. CD62L (L-selectin), a marker of naïve and central memory T cells. MHCII, involved in antigen presentation and expressed on antigen-presenting cells. CD44, an adhesion molecule involved in cell activation and memory T cells. CD11B, expressed on monocytes, macrophages, and neutrophils. FOXP3, a transcription factor identifying Treg cells. TCRβ, a part of the T cell receptor complex, crucial for antigen recognition by T cells.

**Fig. S7.**
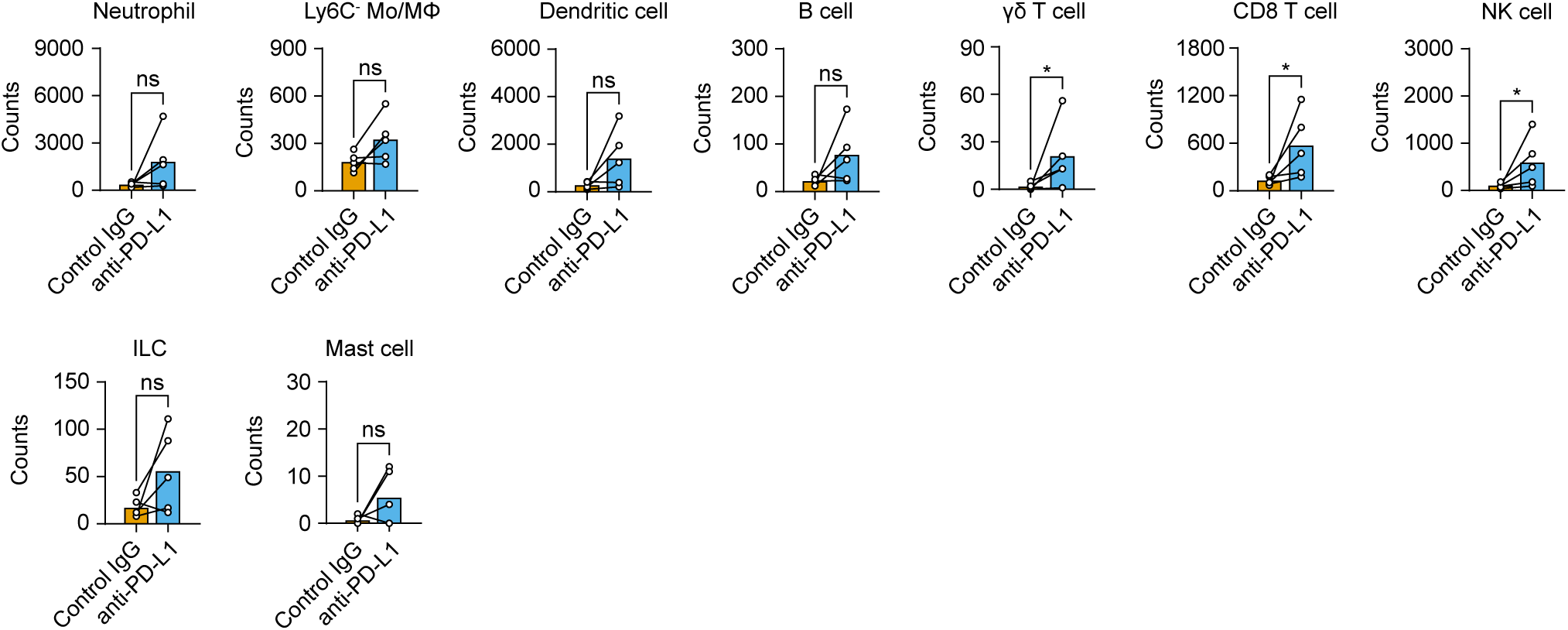
Intracortical anti-PD-L1 injection increases the count of brain infiltrating cells in 5xFAD mice. Counts of brain-infiltrated immune cells in the control IgG- and anti-PD-L1-injected cortex one day after a single intracortical injection (*n* = 5 mice, Wilcoxon signed-rank test; neutrophil; *p* = 0.138, Ly6C-Mo/MΦ; *p* = 0.104, dendritic cell; *p* = 0.080, B cell; *p* = 0.138, γδ T cell; \**p* = 0.043, CD8 T cell; \**p* = 0.043, NK cell; \**p* = 0.043, ILC; *p* = 0.138, mast cell; *p* = 0.144, mean ± s.e.m., ns, not significant).

**Fig. S8.**
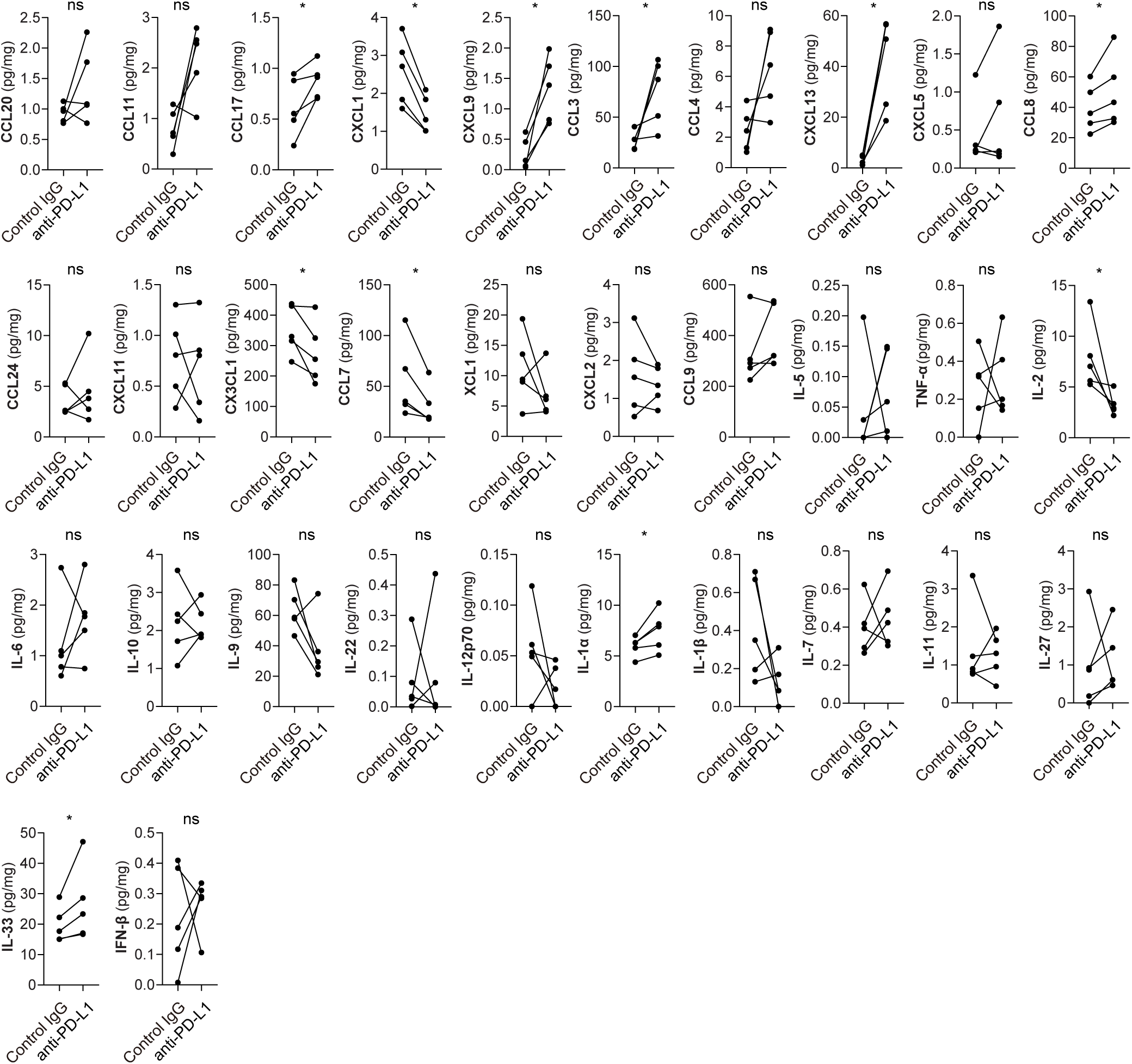
Cytokine and chemokine levels one day after a single intracortical injection of control IgG and anti-PD-L1. The quantification of cytokines and chemokines one day after a single intracortical injection of control IgG and anti-PD-L1 (*n* = 5 mice, Wilcoxon signed-rank test; CCL20; *p* = 0.225, CCL11; *p* = 0.080, CCL17; \**p* = 0.043, CXCL1; \**p* = 0.043, CXCL9; \**p* = 0.043, CCL3; \**p* = 0.043, CCL4; *p* = 0.080, CXCL13; \**p* = 0.043, CXCL5; *p* = 0.500, CCL8; \**p* = 0.043, CCL24; *p* = 0.050, CXCL11; *p* = 0.892, CX3CL1; \**p* = 0.043, CCL7; \**p* = 0.043, XCL1; *p* = 0.345, CXCL2; *p* = 0.345, CCL9; *p* = 0.225, IL-5; *p* = 0.500, TNF-α; *p* = 0.893, IL-2; \**p* = 0.043, IL-6; *p* = 0.345, IL-10; *p* = 0.893, IL-9; *p* = 0.080, IL-22; *p* = 0.893, IL-12p70; *p* = 0.225, IL-1α; \**p* = 0.043, IL-1β; *p* = 0.225, IL-7; *p* = 0.686, IL-11; *p* = 0.893, IL-27; *p* = 0.500, IL-33; \**p* = 0.043, IFN-β; *p* = 0.686, mean ± s.e.m., ns, not significant).

**Fig. S9.**
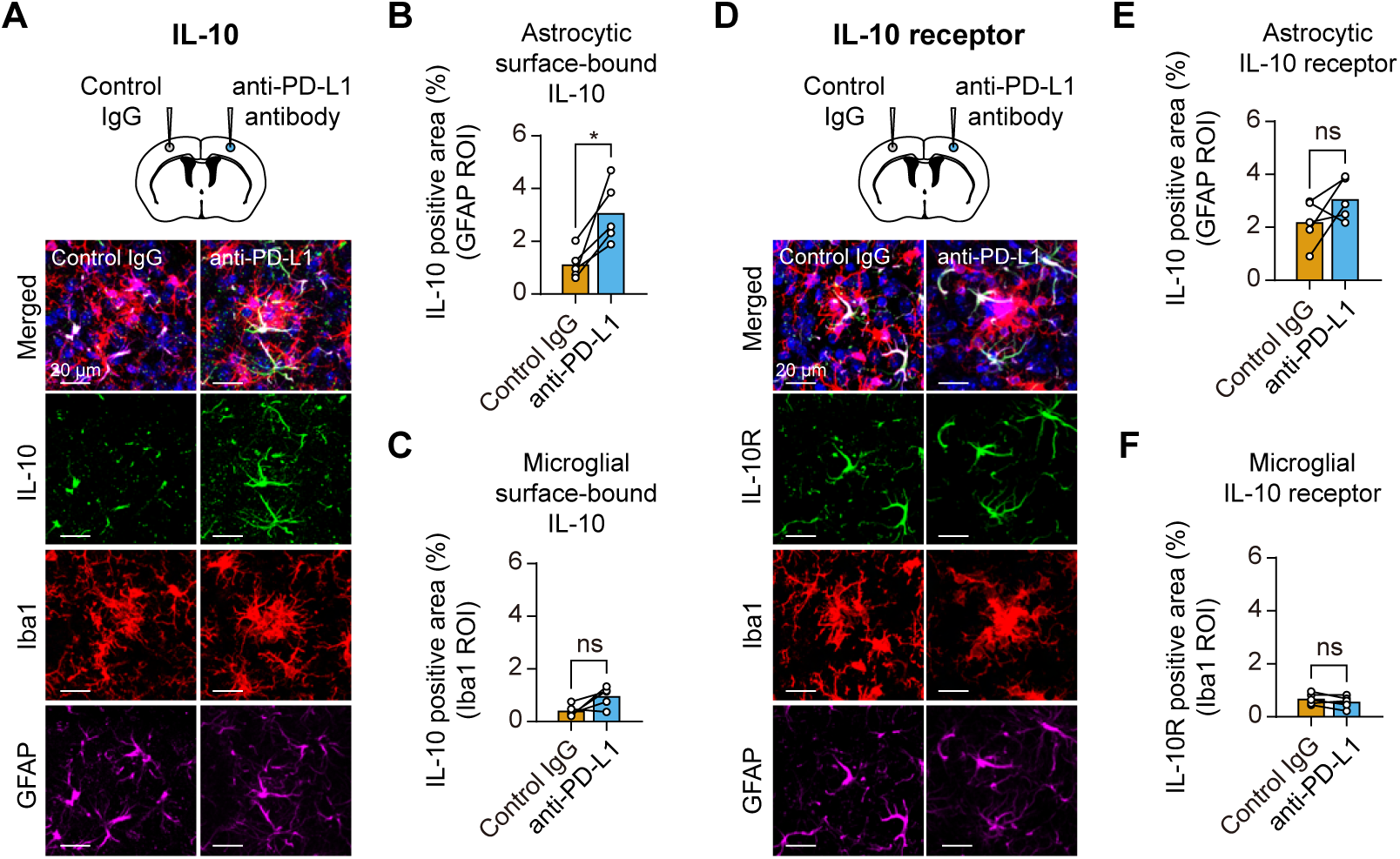
Blockade of PD-1/PD-L1 signaling increases astrocytic surface-bound IL-10 in the brains of 5xFAD mice. (**A**) The experimental scheme and representative IL-10 immunofluorescence images seven days after a single intracortical injection (DAPI, blue; IL-10, green; GFAP, magenta; Iba1, red). (**B**) The quantitation of the co-localized area of IL-10 and GFAP (*n* = 5 mice, Wilcoxon signed-rank test; \**p* = 0.043). (**C**) The quantitation of the co-localized area of IL-10 and Iba1 (*n* = 5 mice, Wilcoxon signed-rank test; *p* = 0.080, ns, not significant). (**D**) The experimental scheme and representative IL-10R immunofluorescence images seven days after a single intracortical injection (DAPI, blue; IL-10R, green; GFAP, magenta; Iba1, red). (**E**) The quantitation of the colocalized areas of IL-10 and GFAP (*n* = 5 mice, Wilcoxon signed-rank test; *p* = 0.138, ns, not significant). (**F**) The quantitation of the colocalized area of IL-10 and Iba1 (*n* = 5 mice, Wilcoxon signed-rank test; *p* = 0.225, ns, not significant).

**Fig. S10.**
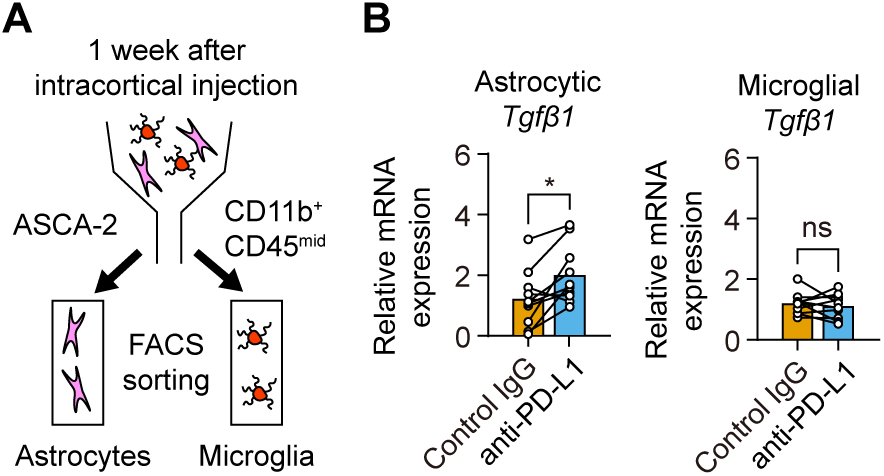
Blockade of PD-1/PD-L1 signaling rescues P2RY12 expression and increases microglial process convergence in response to local brain injury in 5xFAD mice. (**A**) The experimental scheme of the FACS sorting of astrocytes and microglia seven days after a single intracortical injection of control IgG and anti-PD-L1 in 8-month-old 5xFAD mice. (**B**) The relative astrocytic and microglial mRNA levels of *Tgfβ1* (n = 10 mice; Wilcoxon signed-rank test; astrocytes; \**p* = 0.017, microglia; *p* = 0.575, ns, not significant).

**Fig. S11.**
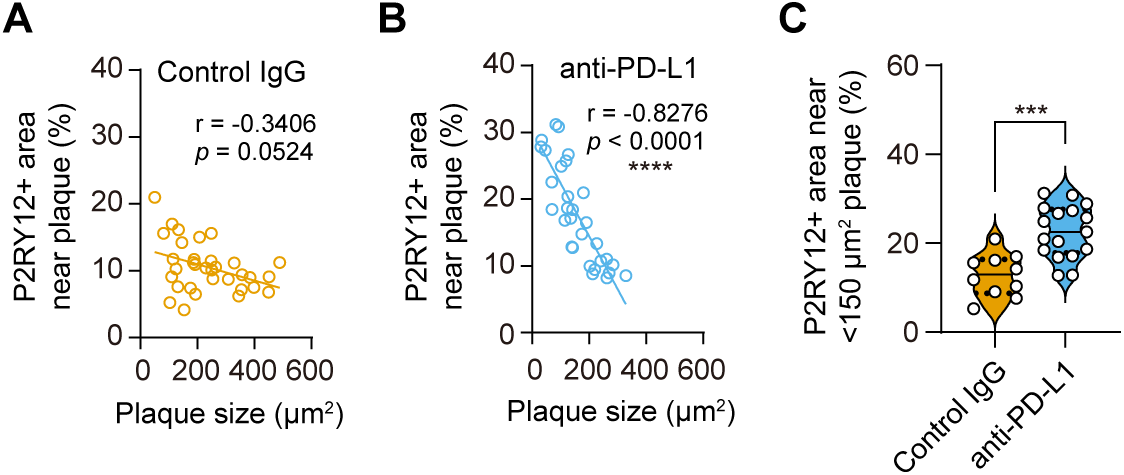
Intracortical injection of anti-PD-L1 in 5xFAD mice increased the P2RY12-positive area in plaque-associated microglia along with the reduction in Aβ plaques. (**A**) The relationship of the P2RY12 positive area with the plaque size in the control IgG-treated area in the 5xFAD mouse (207 plaques, Spearman’s r; -0.3406, *p* = 0.0524). (**B**) The relationship of the P2RY12 positive area with the plaque size in the anti-PD-L1-treated area in the 5xFAD mouse (93 plaques, Spearman’s r; -0.8276, \*\*\*\**p* < 0.0001). (**C**) The quantification of the P2RY12-positive microglial area surrounding the Aβ plaques, each smaller than 150 μm^2^ (Control IgG; 10 plaques, anti-PD-L1; 17 plaques, Mann-Whitney U test; \*\*\**p* = 0.0003).

